# Arabidopsis Myosin XIK Interacts with the Exocyst Complex to Facilitate Vesicle Tethering during Exocytosis

**DOI:** 10.1101/2020.08.18.255984

**Authors:** Weiwei Zhang, Lei Huang, Chunhua Zhang, Christopher J. Staiger

**Author notes:** The author responsible for distribution of materials integral to the findings presented in this article in accordance with the policy described in the Instructions for Authors is: Christopher J. Staiger.

## Abstract

Myosin motors are essential players in secretory vesicle trafficking and exocytosis in yeast and mammalian cells; however, similar roles in plants remain a matter for debate, at least for diffusely-growing cells. Here, we demonstrate that Arabidopsis (*Arabidopsis thaliana*) myosin XIK, via its globular tail domain (GTD), participates in the vesicle tethering step of exocytosis through direct interactions with the exocyst complex. Specifically, myosin XIK GTD bound directly to the SEC5B subunit of exocyst in vitro and functional fluorescently-tagged XIK colocalized with multiple exocyst subunits at plasma membrane (PM)-associated stationary foci. Moreover, genetic and pharmacological inhibition of myosin XI activity reduced the frequency and lifetime of stationary exocyst complexes at the PM. By tracking single exocytosis events of cellulose synthase (CESA) complexes (CSCs) with high spatiotemporal resolution imaging and pair-wise colocalization analysis of myosin XIK, exocyst subunits and CESA6, we demonstrated that XIK associates with secretory vesicles earlier than exocyst and is required for the recruitment of exocyst to the PM tethering site. This study reveals an important functional role for myosin XI in secretion and provides new insights about the dynamic regulation of exocytosis in plants.

## INTRODUCTION

Exocytosis involves the production and trafficking of secretory vesicles from the Golgi to the plasma membrane (PM) where they are tethered and ultimately fuse to deliver new membrane, extracellular matrix components, and membrane-associated proteins. In plants, secretory trafficking and exocytosis coordinate the delivery of lipids, protein receptors, enzymes, polysaccharides and other molecules that are fundamental for many aspects of plant growth and survival, including cell wall biogenesis, cytokinesis, cell polarity establishment, and response to environmental stresses (Friml, 2010; McFarlane et al., 2014; Robatzek, 2014; Kim and Brandizzi, 2016; Yun and Kwon, 2017; Elliott et al., 2020).

Exocytosis is a precisely choreographed process requiring the spatiotemporal cooperation of a plethora of molecules and protein complexes, including cytoskeleton and motor proteins, Rab-family GTPases, the exocyst tethering complex, and soluble *N*-ethylmaleimide sensitive factor attachment protein receptors (SNAREs) to ensure targeted delivery of cargos to specific PM regions for secretion. The exocyst is a conserved octameric protein complex that mediates the initial tethering of secretory vesicles to the PM before SNARE-mediated membrane docking and fusion (TerBush et al., 1996; Cvrčková et al., 2012; Žárský et al., 2013; Wu and Guo, 2015; Ravikumar et al., 2017). The eight subunits comprising the exocyst complex are SEC3, SEC5, SEC6, SEC8, SEC10, SEC15, EXO70 and EXO84. Although many of the core components in exocytosis are evolutionarily conserved and well characterized in animal and yeast models, our knowledge of the precise mechanisms and molecular players that coordinate secretion in plant cells remains sparse.

A key player in exocytosis in animal and yeast cells is the class V myosin, whose primary role is transport of vesicles and organelles along the actin cytoskeleton (Rudolf et al., 2011; Hammer and Sellers, 2012). Arguably, the best characterized myosin V is budding yeast Myo2p, which plays a vital role in polarized growth through powering the transport of secretory vesicles along actin cables from the mother cell to the growing tip in the bud (Govindan et al., 1995). Myo2p transports secretory vesicles in a cargo receptor-dependent manner through interaction with the Rab GTPase receptor SEC4 via its globular tail domain (GTD; Jin et al., 2011; Santiago-Tirado et al., 2011). SEC4 also recruits the exocyst tethering complex to the vesicle surface by direct association with the SEC15 subunit (Guo et al., 1999). The GTD of Myo2p also binds to SEC15, which is critical for localization of SEC15 to the bud tip and polarized secretion (Jin et al., 2011). Myo2p remains associated with the vesicle after it arrives at the PM tethering site until a few seconds before the vesicle fully fuses with the PM, and regulates vesicle tethering time through an unknown mechanism that is independent of its interaction with SEC15 (Donovan and Bretscher, 2012, 2015). Similarly, myosin V motors in other species interact with the homologous Rab GTPases or other cargo receptors to facilitate secretory vesicle transport (Jin et al., 2011; Lindsay et al., 2013; Vogel et al., 2015). One recent study demonstrates a role for myosin V in neuronal synapses by tethering synaptic vesicles to PM release sites in an ATPase activity-dependent manner, rather than driving vesicle transport (Maschi et al., 2018). Myosin V also contributes to vesicle tethering through Ca^2+^-dependent interaction with SNARE proteins on synaptic vesicles (Prekeris and Terrian, 1997; Watanabe et al., 2005). In other mammalian cell types, such as endocrine and neuroendocrine cells, myosin V controls the targeted transport and exocytosis of hormones and neuronal peptides by capturing or tethering secretory vesicles in a cortical actin meshwork near the exocytosis site to prevent premature fusion (Rudolf et al., 2011).

Myosin V-like motors are widely present in plants and are grouped into myosin VIII and XI families (Reddy and Day, 2001). Despite the well characterized roles of myosin V in secretory vesicle transport and exocytosis, there is a paucity of information about similar roles for plant myosin. Conventional wisdom maintains that the major role of plant myosin XI is to power cytoplasmic streaming and long-distance transport of organelles and vesicles (Avisar et al., 2012; Tominaga and Ito, 2015; Ueda et al., 2015). In *Arabidopsis thaliana*, the myosin XI family comprises 13 members, with XIK, XI1 and XI2 among the most highly expressed isoforms that function redundantly in driving intracellular motility and thereby contributes to rapid cell growth and expansion (Prokhnevsky et al., 2008; Peremyslov et al., 2010; Ueda et al., 2010; Haraguchi et al., 2018). Further, myosin XIK is the primary myosin responsible for cytoplasmic streaming and organelle motility (Avisar et al., 2012).

In addition to Rabs and the exocyst complex, numerous cargo receptors and myosin V-interacting proteins have been identified in yeast and animal cells. By comparison, the major myosin XI-binding proteins identified are plant-specific MyoB family proteins (Peremyslov et al., 2013; Kurth et al., 2017). Surprisingly, none of the MyoBs in Arabidopsis colocalize with large organelles or secretory vesicles, but instead are associated with a specific type of endomembrane compartment that moves rapidly along actin cables to drive cytoplasmic streaming; consequently, an indirect model of myosin XI powering transport of organelles in plant cells has been proposed (Peremyslov et al., 2013; Kurth et al., 2017; Nebenführ and Dixit, 2018). Additional myosin XI interactors include MadA/B family proteins (Kurth et al., 2017), DECAPPING PROTEIN1 (Steffens et al., 2014), and WIT1/2 (Tamura et al., 2013), none of which have been implicated in secretory trafficking or directly connect myosin XI to secretory vesicles. Nevertheless, there is emerging evidence that myosin XI may play a role in exocytosis, at least in tip-growing cells (Orr et al., 2020). In the moss *Physcomitrella patens*, myosin XI interacts with a RabE GTPase that is homologous to yeast Sec4; disruption of this binding results in unpolarized growth (Orr et al., 2019). Another study in tobacco pollen tubes showed that the MyoB family protein RISAP interacts with the RAC/ROP GTPase RAC5 and is proposed to mediate secretory trafficking during pollen tube tip growth (Stephan et al., 2014). However, whether a tripartite complex of myosin XI tail domain, exocyst subunits, and Rab functions in plants to mediate secretory vesicle tethering remains to be established.

Delivery of polysaccharides and proteins to construct the cell wall provides a facile system to dissect exocytic trafficking in plant cells. Cellulose, the primary component of the cell wall, is synthesized at the cell surface by cellulose synthase (CESA) complexes (CSCs) which rely on endomembrane trafficking and exocytosis for delivery to the PM (Bashline et al., 2014; McFarlane et al., 2014). Recently, the plant exocyst complex has been implicated in mediating delivery of CSCs to the PM in primary (Zhu et al., 2018) and secondary cell wall deposition (Vukašinović et al., 2017). The exocyst complex associates with CSCs for a few seconds during the initial static phase after the vesicle has arrived at the PM, consistent with a role in tethering of CESA compartments to the PM (Zhu et al., 2018). In previous work, we showed that myosin XI is also a key player in delivery of CSCs to the PM (Zhang et al., 2019). Myosin XI regulates the rate of CSC delivery to the PM, arrives at the PM along with putative secretory vesicles and associates transiently at the docking site, and facilitates vesicle tethering or fusion. The detailed molecular mechanisms for myosin XI in vesicle tethering/fusion, however, remain unclear and whether interactions with other players, such as the exocyst complex, are required merits further investigation.

Here, we report that myosin XIK is the primary myosin isoform mediating CSC delivery, vesicle tethering, and exocytosis in Arabidopsis. Moreover, yeast two-hybrid and in vitro pull-down assays revealed a direct interaction between myosin XIK GTD and the SEC5B subunit of exocyst complex. By combining genetic and pharmacological approaches with quantitative live-cell imaging of Arabidopsis lines expressing combinations of fluorescent reporters for CESA6, myosin XIK, or exocyst subunits, we showed that myosin XI regulates exocyst dynamics at the PM and is required for the localization of exocyst at vesicle tethering sites during CSC delivery at the PM. Collectively, these data demonstrate the exocyst complex is a new interactor of myosin XI and we propose a novel role for myosin XI in regulating exocytosis in plants.

## RESULTS

### Myosin XIK is a Major Isoform Involved in Cellulose Biogenesis and CESA Trafficking

Using the trafficking of CSCs as a model experimental system, Arabidopsis myosins XI were shown to play a role in vesicle tethering and/or fusion at the PM (Zhang et al., 2019). However, it remains unclear how individual myosin isoforms are involved in this process and whether there is one isoform that plays a predominant role. An Arabidopsis *myosin xi1 xi2 xik* triple-knockout mutant (*xi3KO*) has reduced cellulose levels and a significantly lower CSC delivery rate (Zhang et al., 2019). Myosin XI1, XI2, and XIK isoforms are among the most highly-expressed myosins in Arabidopsis somatic cells, with XIK known to be the primary isoform responsible for cytoplasmic streaming and organelle transport (Peremyslov et al., 2008; Prokhnevsky et al., 2008; Ueda et al., 2010; Avisar et al., 2012; Haraguchi et al., 2018). It is plausible that XIK is also the major isoform involved in CESA trafficking. To test this possibility, we screened *xik-2, xi1,* and *xi2* single gene knockout mutants (Peremyslov et al., 2010) for cellulose biosynthesis and CESA trafficking defects. Measurement of cellulose content was conducted using the trifluoroacetic acid (TFA) and acetic-nitric (AN) methods (Zhang et al., 2019). The results showed that the greatest decrease in total and crystalline cellulose levels occurred in *xik-2*, with the extent of reduction comparable to that in the *xi3KO* mutant, whereas only a slight reduction was detected in *xi2* and no obvious effect was observed in *xi1* (Figures 1A and 1B). To confirm the phenotype found in *xik-2*, a second knockout mutant allele *xik-1* (Ojangu et al., 2007) was analyzed which showed similar reduction in cellulose content when compared with *xik-2* (Figures 1A and 1B).

**Figure 1.**
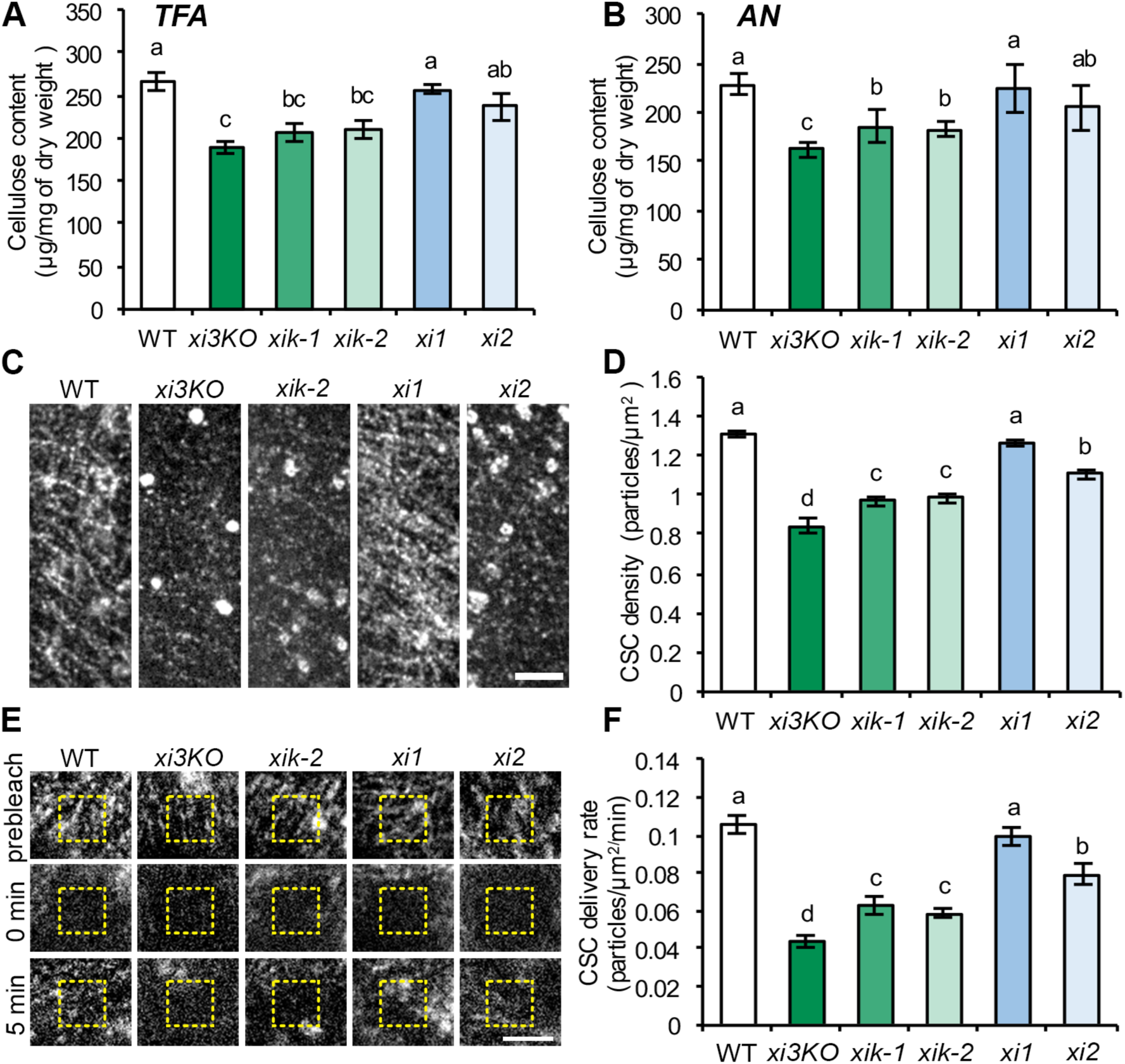
Myosin XIK Is the Major Myosin Isoform Involved in Cellulose Biogenesis and CESA Trafficking. (**A, B**) XIK contributes to cellulose production. Ethanol-insoluble cell wall material (CWM) was prepared from 5-d-old etiolated hypocotyls of wild-type (WT) seedlings, *myosin xi3KO*, *xik-1*, *xik-2*, *xi1,* and *xi2* mutants. The non-cellulosic component of CWM was hydrolyzed with 2 M trifluoroacetic acid (TFA; **A**) for total cellulose determination, or with acetic nitric reagent (AN; **B**) for crystalline cellulose determination. Cellulose content was significantly reduced in *xi3KO*, *xik-1*, and *xik-2* mutants compared to that in WT, *xi1,* and *xi2* mutants. Values given are means ± SE (n = 4; One-way ANOVA with Tukey’s post hoc test, letters [a-c] denote samples/groups that show statistically significant differences from other groups, P < 0.05). (**C, D**) XIK is necessary for the abundance of CSCs at the PM. Representative single-frame images show the plasma membrane (PM) of hypocotyl epidermal cells expressing YFP-CESA6 imaged with spinning disk confocal microscopy (**C**). Bar = 5 µm. Quantitative analysis shows that the density of CSC at the PM was significantly reduced in *xi3KO*, *xik-1*, *xik-2,* and *xi2*, but not in *xi1* (**D**). Values given are means ± SE (n > 60 cells from 12 hypocotyls per genotype; One-way ANOVA with Tukey’s post hoc test, letters [a-d] denote samples/groups that show statistically significant differences from other groups, P < 0.05). (**E, F**) Loss of XIK reduces the rate of delivery of CSCs to the PM. Representative single-frame images of PM-localized CSC particle recovery after photobleaching. A region of interest at the PM was photobleached and the number of newly-delivered CSCs were counted in a subarea within the region (yellow dashed box, **E**). Bar = 5 µm. The rate of delivery of CSCs to the PM was calculated from the total number of newly-delivered CSCs during the initial 5 min of recovery divided by the measured area and time (**F**). The CSC delivery rate was significantly inhibited in *xi3KO*, *xik-1*, *xik-2,* and *xi2*, but not in *xi1*. Values given are means ± SE (n = 9–12 cells per genotype; One-way ANOVA with Tukey’s post hoc test, letters [a-d] denote samples/groups that show statistically significant differences from other groups, P < 0.05).

To test whether a single myosin isoform makes a major contribution to exocytosis, we analyzed CESA trafficking phenotypes by quantitative live cell imaging with spinning disk confocal microscopy (SDCM) in 3-d-old etiolated hypocotyl epidermal cells using *xi1*, *xi2, xik-1,* and *xik-2* single mutants expressing YFP-CESA6 in the *prc1-1* homozygous mutant background. Among these lines, *xi1*, *xi2,* and *xik-2* single mutants were recovered from the same cross that resulted in the previously characterized *xi3KO* YFP-CESA6 *prc1-1* line (Zhang et al., 2019). Similar to the previous report (Zhang et al., 2019), the *xi3KO* YFP-CESA6 *prc1-1* line exhibited significantly decreased density (35%) of PM-localized CSCs and a 59% reduction of CSC delivery rate to the PM compared with wild-type siblings (Figures 1C to 1F). Analysis of myosin single mutants revealed that *xik-1* and *xik-2* showed the most severe disruption of CSC trafficking, with CSC density decreased by ∼25% and the delivery rate reduced by ∼40% compared with wild-type siblings, whereas *xi2* had a slight reduction and *xi1* showed no significant reduction in either density and delivery rate assays (Figures 1C to 1F). These results indicate that XIK is a major player in delivery of CSCs to the PM and makes a substantial contribution to cellulose production. Hereafter, we use the two *xik* single mutant alleles to further characterize the role of Myosin XI in delivery, tethering, and fusion of vesicles at the PM.

### Myosin XIK Mediates the Exocytosis of CSCs

The *xi3KO* mutant has vesicle tethering or fusion defects as well as an abnormal accumulation of CESA-containing compartments in the cortical cytoplasm, likely resulting from inefficient exocytosis (Zhang et al., 2019). To test the role of XIK in these processes, we initially measured the abundance of cortical and subcortical CESA compartments in *xik-1* and *xik-2*. Using previously described methods (Sampathkumar et al., 2013; Zhang et al., 2019), we imaged epidermal cells from 3-d-old etiolated hypocotyls with SDCM and combined the optical sections into cortical (0 to 0.4 µm below the PM) and subcortical (0.6 to 1 µm below the PM) cytoplasm. The reason for segmenting the cytoplasm into two regions is because if myosin participates in vesicle exocytosis, an abnormal CESA compartment population is more likely to be detected in the cortex close to the PM rather than in the subcortical region. Similar to observations in *xi3KO* (Zhang et al., 2019), an increased population of CESA compartments was detected only in the cortical cytoplasm but not in the subcortical region of *xik-1* and *xik-2* cells compared with that in wild-type cells (Figures 2A and 2B).

**Figure 2.**
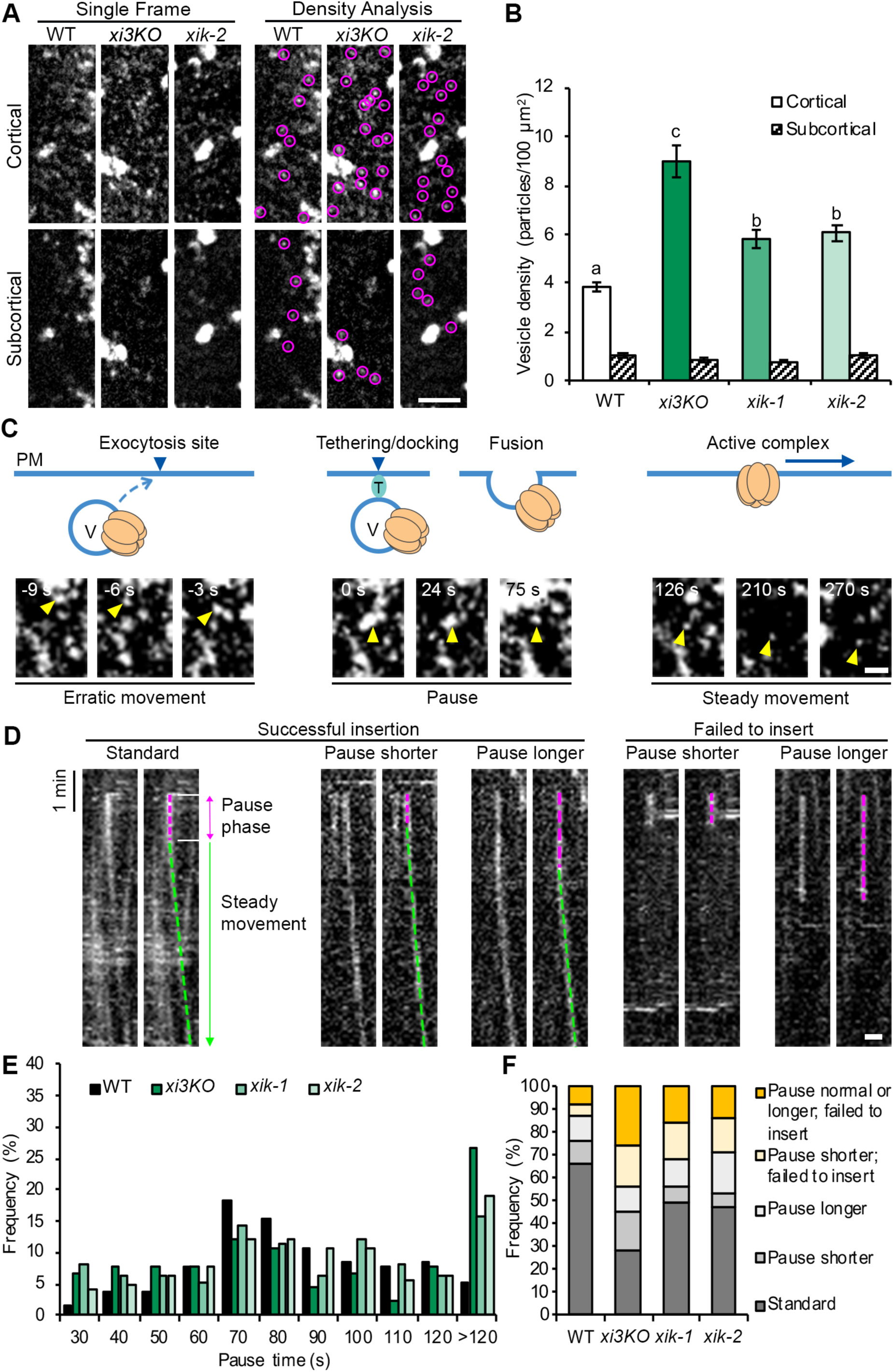
Myosin XIK Is Involved in Exocytosis of CSCs. (**A, B**) Loss of XIK results in increased abundance of cortical vesicles containing CESA6. Representative single images taken at cortical and subcortical focal planes in hypocotyl epidermal cells show cytoplasmic CESA compartments (magenta circle) in WT, *xi3KO,* and *xik* (**A**). Bar = 5 µm. Quantitative analysis of vesicle density shows that the number of CESA compartments was increased significantly in the cortical but not in the subcortical cytoplasm for *xi3KO*, *xik-1*, and *xik-2* compared to WT siblings (**B**). Values given are means ± SE (n > 25 cells from 12 seedlings for each genotype; One-way ANOVA with Tukey’s post hoc test, letters [a-c] denote samples/groups that show statistically significant differences from other groups, P < 0.05). (**C-F**) XIK is necessary for CSC tethering and fusion at the PM. Representative images show a typical CSC insertion event at the PM (**C**). A CESA particle (yellow arrowhead) arriving in the cortex initially undergoes erratic motility, which likely represents a delivery vesicle (V) that is transported to an exocytosis site. The particle then pauses (marked as 0 s) and exhibits a static or pause phase for ∼80 s in a fixed position, which likely corresponds to tethering, docking and fusion of the delivery compartment to the PM. After the CSC particle is inserted, it shows steady movement in the PM as an active complex. T: tethering proteins. Bar = 1 µm. (**D**) Representative kymographs show five categories of insertion events from left to right: standard insertion with normal pause time; insertion that has a shorter pause time; insertion that has a longer pause time; a shorter pause time and failure to insert; and, a longer pause time and failure to insert. Pause phases are marked with magenta dashed lines and steady movement phases are marked with green dashed lines. Bar = 1 µm. (**E**) Distribution of pause times during CSC insertion at the PM in WT and mutant hypocotyl epidermal cells. (n ≥ 10 cells from 7–10 seedlings for each genotype; a total of 119, 132, 159, and 141 events were measured in WT, *xi3KO*, *xik-1,* and *xik-2* cells, respectively). (**F**) The proportion of five types of insertion events described in (**D**) in WT and mutant hypocotyl epidermal cells. A shorter or longer pause time was defined as the mean value (84 ± 30 s) of particle pause time in WT minus (< 54 s) or plus one standard deviation (> 114 s), respectively.

To further investigate the role of XIK in vesicle secretion and test whether exocytosis defects result from loss of XIK, we performed a single CSC insertion assay using high-resolution spatiotemporal imaging with SDCM. A single CSC insertion event typically undergoes three phases of dynamic movement: a transient erratic phase, representing a vesicle approaching its PM destination for fusion; a pause phase at the PM that lasts for 1 to 2 min, representing vesicle tethering, docking and fusion; and a steady movement phase, indicative of an active cellulose-producing complex moving along a linear trajectory in the PM (Figure 2C; Supplemental Movie 1; Gutierrez et al., 2009; Zhang et al., 2019). By tracking the dynamics of new CSC insertion events, we previously showed that they can be categorized into five groups: (1) a standard event with a pause time for 1 to 2 min followed by a steady movement phase following successful CSC insertion; (2) a successful insertion event with a shorter pause time (shorter than the mean pause time in wild type minus one standard deviation); (3) a successful insertion with a longer pause time (longer than the mean pause time in wild type plus one standard deviation); (4) a failed insertion with only a shorter pause phase and no steady movement phase; (5) a failed insertion with a normal or longer pause time (Figure 2D; Zhang et al., 2019). Through analysis of kymographs, we quantified pause times and the frequency of each type of insertion event in the myosin mutant lines. In wild type, the average pause time was 84 ± 29 s (mean ± SD, n = 131 events), similar to the previously reported value of 81 ± 27 s (Zhang et al., 2019). In contrast, in the two *xik* alleles and the *xi3KO* mutant, a number of events exhibited an abnormal pause time, with 15–27% of the events showing pause times longer than 120 s compared with that of 5% in wild-type cells (Figure 2E). In addition, the population with shorter pause times of 15–50 s also increased from 9% in wild type to 16–22% in myosin single or triple mutants (Figure 2E). Moreover, the majority of events with abnormal pause times in myosin mutants also failed to insert functional CSCs into the PM (Figure 2F). The percentage of total failed insertion events increased from 13% in wild type to 44, 32, and 29% in *xi3KO*, *xik-1,* and *xik-2*, respectively (Figure 2F). As the CSC pause phase is shown to be associated with vesicle tethering and fusion (Zhu et al., 2018), the altered CSC pause time and increased frequency of failed insertion events in *xik* mutants indicate that XIK plays a role in the tethering or fusion step during the exocytosis of CSC-containing vesicles at the PM.

Consistent with a role for myosin XI in vesicle tethering, we showed previously that functional YFP-tagged XIK displayed transient colocalization with tdTomato-CESA6 during the first 3 s of the pause phase during CSC insertion at the PM (Zhang et al., 2019). Given that tethering is likely to last for ∼10 s at the beginning of the pause phase as indicated by the colocalization of exocyst tethering complex with CSCs (Zhu et al., 2018), we speculate that myosin and exocyst may cooperate to achieve vesicle tethering.

### Myosin XIK Interacts Directly with Exocyst Subunits

A direct interaction between the GTD of Myo2p and the exocyst subunit SEC15 has been demonstrated in budding yeast (Jin et al., 2011). To explore this relationship in plants, we tested for direct interactions using a yeast two-hybrid assay with Arabidopsis exocyst subunits as prey and the GTD of XIK as bait. Through a screen of members from all eight subunits of the exocyst complex, SEC5B strongly interacted with XIK GTD and EXO84A showed a weak interaction (Figure 3A). The direct interaction of XIK GTD with SEC5B was further verified with an in vitro pull-down assay (Figure 3B). Recombinant SEC5B protein cosedimented with purified GST-tagged XIK GTD but not with purified GST alone (Figure 3B). The direct interaction of XIK with exocyst subunits supports our finding that XIK plays a role in vesicle tethering in plants.

**Figure 3.**
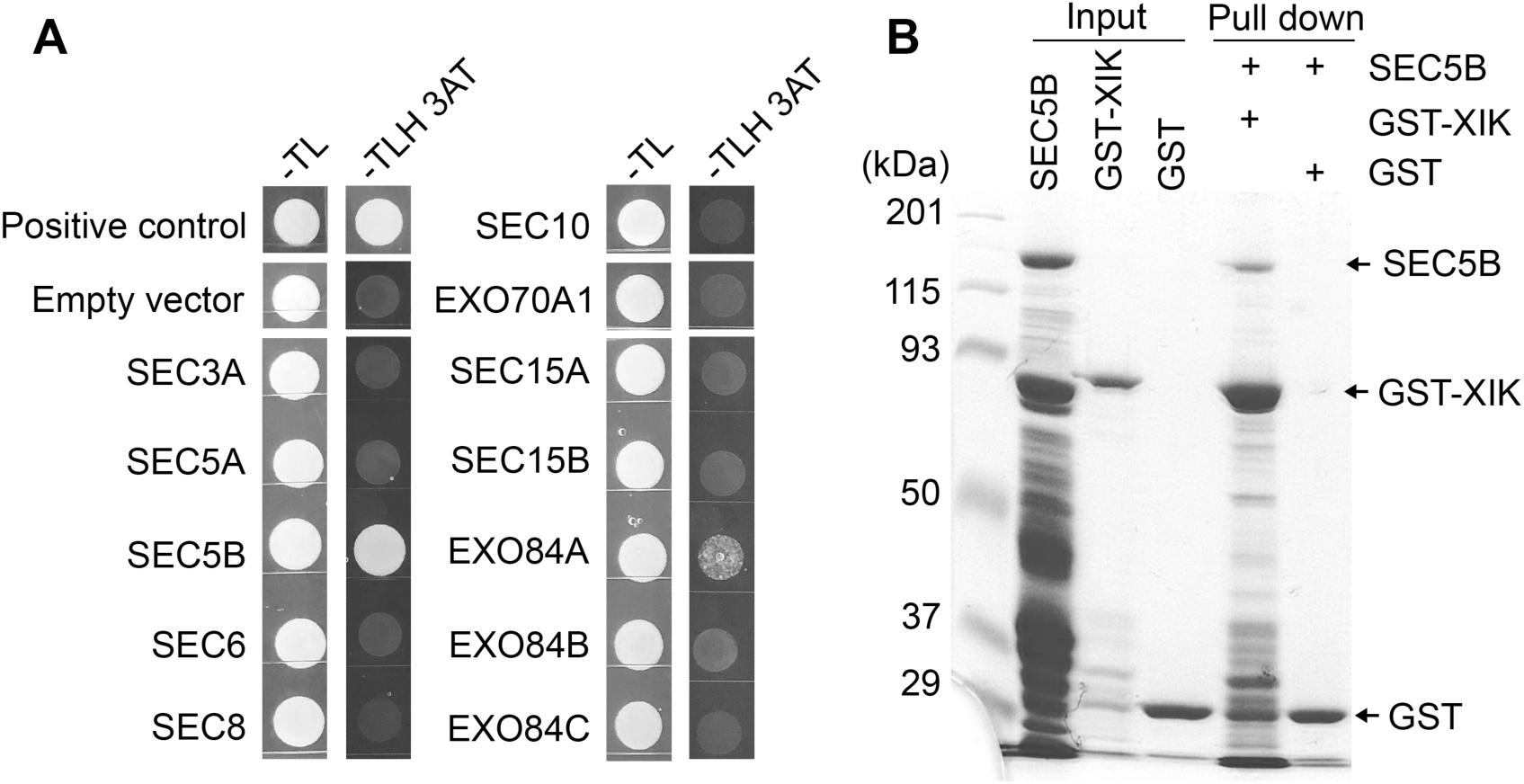
The Globular Tail Domain (GTD) of Myosin XIK Interacts Directly with Exocyst Subunits. **(A)** Yeast two-hybrid with XIK GTD as bait and exocyst subunits as prey. The co-transformed colonies were grown on SD-Trp-Leu (-TL) plates or SD-Trp-Leu-His plates supplemented with 4 mM 3-amino-1,2,4-triazole (-TLH 3AT). Growth of colonies shown on -TLH 3AT plates when SEC5B and Exo84A were used as prey indicates interactions with XIK GTD. **(B)** Protein pull-down assay shows interaction between XIK GTD and SEC5B in vitro. Purified, recombinant His6-tagged SEC5B cosedimented with purified XIK GTD fused to a GST tag (GST-XIK) but not with the purified GST protein alone.

### Myosin XIK Regulates the Dynamic Behavior of Exocyst Foci at the PM

Given the direct interaction between XIK and the exocyst complex, we tested whether the two are functionally associated in plants. Dynamic analysis of several exocyst subunits in Arabidopsis has shown that those subunits localize in discrete foci at the PM (Fendrych et al., 2013; Zhang et al., 2013). We crossed a functional EXO70A1-GFP reporter (Fendrych et al., 2010) into *xik-1* and *xik-2* and recovered homozygous mutant lines as well as wild-type siblings expressing EXO70A1-GFP. In addition to the genetic mutation of XIK, we also applied acute drug treatments with the myosin inhibitor pentabromopseudilin (PBP), which potently inhibits XIK-YFP motility in Arabidopsis cells (Zhang et al., 2019). Hypocotyl epidermal cells were imaged with high resolution variable-angle epifluorescence microscopy (VAEM), and the EXO70A1-GFP signal appeared as abundant distinct foci at the PM with a density of ∼1.5 particles/µm^2^ (Figure 4A), similar to previous reports (Fendrych et al., 2013). The overall density and distribution pattern of EXO70A1-GFP foci at the PM were similar in *xik* cells and wild-type cells treated with PBP for 15 min, compared to wild type (Figures 4A and 4C; Supplemental Figure 1), suggesting that XIK is not required for the distribution of exocyst complexes at the PM.

**Figure 4.**
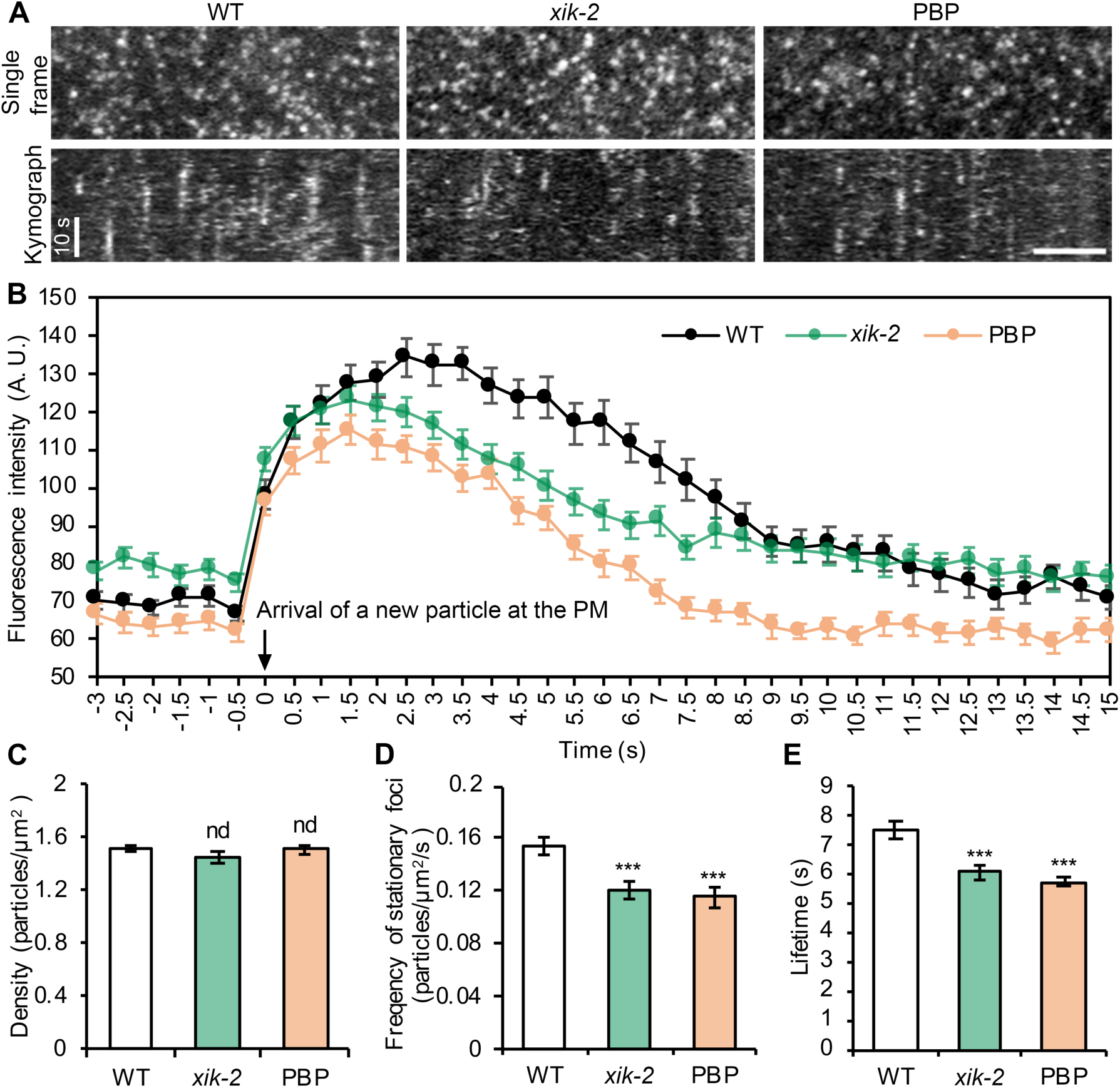
Inhibition of Myosin Activity Alters the Dynamic Behavior of EXO70A1 at the PM. **(A)** Representative single frame images show distribution of exocyst subunit EXO70A1-GFP at the PM in 3-d-old etiolated hypocotyl epidermal cells imaged with variable-angle epifluorescence microscopy. Kymographs reveal the presence of stationary EXO70A1-GFP foci in time-lapse series. There were fewer stationary foci in *xik-2* and cells treated with PBP for 15 min, and their lifetime appeared shorter compared to WT cells. Bar = 5 µm. **(B)** Quantitative analysis of fluorescence intensity for newly-appearing stationary foci of EXO70A1-GFP at the PM. The zero timepoint was defined as the first frame in which a new particle appears. Values given are means ± SE (n = 71, 88 and 78 particles in WT, *xik-2,* and PBP-treated cells, respectively). (**C–E**) Quantitative analysis shows that the density of total EXO70A1-GFP foci remains similar in *xik-2* and PBP-treated cells (**C**), however, the frequency (**D**) and lifetime (**E**) of stationary foci were significantly reduced compared with that in WT cells. Values given are means ± SE (n = 20–30 cells from 10 seedlings per genotype or treatment; for lifetime assay, n = 71, 88 and 78 particles in WT, *xik-2* and PBP-treated cells, respectively; Student’s t test, nd: P > 0.05, ***P < 0.001).

When we tracked the dynamic behavior of the EXO70A1 foci at the PM, two distinct populations were observed: one population exhibited a stationary phase following their appearance at the PM, whereas the other population did not have a pause phase but showed short and rapid diffuse motility before disappearing from the plane of the PM (Supplemental Movie 2). We focused on the stationary population, as it has been suggested that not all exocyst foci tether a secretory vesicle and foci with a stationary phase at the PM are more likely to represent real exocytic events (Fendrych et al., 2013). A stationary exocyst particle was defined by a straight vertical line in kymographs prepared from time-lapse series (Figure 4A) that could be tracked for at least 5 frames (> 2 s). Quantification of the number of newly-appeared stationary foci over time showed that the frequency of these foci at the PM was significantly decreased by ∼25% in *xik* and PBP-treated cells compared with wild-type cells (Figure 4D; Supplemental Figure 1; Supplemental Movie 2). As the stationary foci are likely to be linked to vesicle tethering/exocytosis events (Fendrych et al., 2013), the results suggest that even though XIK activity is not required for overall exocyst localization at the PM, it may be necessary for the PM targeting of a subpopulation that is responsible for vesicle tethering.

We next analyzed the fluorescence intensity profiles of newly-appeared stationary foci at the PM. A small fixed region of interest (ROI) that covered the centroid of an exocyst particle was analyzed over a time course, from a few seconds before the appearance of the foci to a few seconds after the foci fully disappeared. By quantifying the average fluorescence intensity from multiple events, the results showed that in wild-type cells, after the appearance of EXO70A1 particles at 0 s, the average fluorescence intensity quickly reached a high level and lasted for 6–7 s before it decreased to background level at around ∼ 9 s (Figure 4B). However, in *xik-2* and PBP-treated cells, the foci displayed an apparently shorter lifetime, with an average signal that peaked and then disappeared about 1–2 s earlier than in wild-type cells (Figure 4B). The average lifetime of EXO70A1 foci in wild type was 7.5 s, whereas in *xik-2* and PBP-treated cells, the lifetime was reduced to 6.1 s and 5.7 s, respectively (Figure 4E). A similar reduction of lifetime of EXO70A1 foci was detected in the *xik-1* mutant when compared with wild-type siblings (Supplemental Figure 1). These results indicate that XIK is required for maintaining normal dynamic properties of exocyst at the PM and disruption of XIK activity results in a shorter exocyst tethering time.

Given that SEC5B directly interacts with myosin XIK GTD in vitro, we generated a GFP-SEC5B reporter line under the control of its native promoter and tested whether the dynamic behavior of SEC5B was also altered upon disruption of myosin XI activity. At the PM plane of hypocotyl epidermal cells, GFP-SEC5B appeared as dense puncta that show rapid dynamic behavior similar to that of EXO70A1-GFP foci and previous reports (Supplemental Figure 2; Fendrych et al., 2013; Zhu et al., 2018). Similar to EXO70A1 foci, we did not detect apparent alteration in overall density or distribution pattern of SEC5B foci at the PM in *xik-2* or after acute PBP treatment; however, the particle dynamic analysis showed that there was a reduced frequency and a shorter lifetime of stationary SEC5B foci in myosin-deficient cells (Supplemental Figure 2). Furthermore, alteration of PM dynamics was observed with a third exocyst subunit marker, SEC6-GFP (Fendrych et al., 2013), when myosin activity was inhibited (Supplemental Figure 3).

Since myosins are actin-based motors, it is likely that the cortical actin cytoskeleton is also involved in regulating exocyst dynamics at the PM. Fendrych et al. (2013) report that a 10-min treatment with the actin polymerization inhibitor latrunculin B (LatB) did not alter exocyst subunit density and distribution at the PM, whereas prolonged treatment (1 h) resulted in aggregation and uneven distribution of exocyst subunits at the PM. Similarly, we observed that a 15-min treatment with 10 μM LatB did not change the overall density or distribution of exocyst foci at the PM, however, the stationary population was affected similarly to that in *xik* or PBP-treated cells, with significantly reduced frequency and a shorter lifetime compared to that in untreated cells (Supplemental Figure 3). These results suggest that both myosin XI and cortical actin are required for exocyst dynamics at the PM.

Collectively, these data indicate that the frequency and lifetime of stationary exocyst foci at the PM are dependent on myosin XI and actin function, thereby supporting the functional association between myosin and exocyst in plant cells.

### Myosin XIK Transiently Colocalizes with Exocyst Foci Near the PM

Given the apparent functional connection between myosin XI and stationary exocyst foci at the PM, we tested whether XIK and these foci colocalize. A transgenic line co-expressing XIK-mCherry (Peremyslov et al., 2013) and GFP-SEC5B under the control of their native promoters was generated and imaged by time-lapse SDCM. The co-expression line was in the *xik-2* mutant background to avoid overexpression of XIK protein which may lead to excess cytoplasmic signal. Similar to previous reports, XIK-mCherry was mostly localized in the cell cortex, with a majority decorating an unknown type of endomembrane compartment that rapidly translocates along actin filaments to drive cytoplasmic streaming, and the rest was present as dynamic patches or diffuse signal in the cytoplasm (Supplemental Movie 3; Peremyslov et al., 2012; Zhang et al., 2019). We previously reported that XIK-YFP cytoplasmic patches transiently and specifically colocalized with newly-arrived CSCs at the site of insertion at the PM (Zhang et al., 2019). Here, we investigated whether there was also specific association of the cytoplasmic XIK signal with newly-appeared SEC5B foci at the PM. Dual-channel time-lapse imaging was performed with SDCM at 1-s intervals and only newly-arrived SEC5B foci in the GFP channel which showed a stationary phase for at least 5 frames (> 4 s) were tracked. Analysis of the corresponding mCherry channel showed that a cluster of cytoplasmic XIK-mCherry signal was frequently observed to colocalize with a new SEC5B particle during the first few seconds upon its arrival at the PM (Figure 5A; Supplemental Movie 3 and 4). Interestingly, the high frequency of spatiotemporal colocalization was mainly observed in the events when SEC5B had a longer lifetime at the PM (measured as the duration of stationary phase in kymographs). SEC5B foci are reported to have a lifetime of 8–12 s (Zhu et al., 2018), and similarly, the mean lifetime of GFP-SEC5B at the PM was measured to be 10 ± 4 s in this assay (n = 202 particles). We found that 70% of the GFP-SEC5B foci with a lifetime of 8 s or longer showed colocalization with XIK-mCherry at the beginning of stationary phase (88 of 126 particles), whereas the colocalization rate was only 41% when the GFP-SEC5B foci had a lifetime shorter than 8 s (31 of 76 particles). To confirm the specific spatiotemporal colocalization, we quantified the fluorescence intensity profiles of individual stationary GFP-SEC5B foci as well as the same ROIs in the corresponding XIK-mCherry channel from time-lapse series (Figure 5C). The results showed that there was significantly higher fluorescence intensity of XIK-mCherry that appeared 2 s before the arrival of a new SEC5B particle, which then peaked at 0 s and lasted for an average of 4 s after the appearance of the new foci at the PM, compared with the remainder of the time points (Figure 5C).

**Figure 5.**
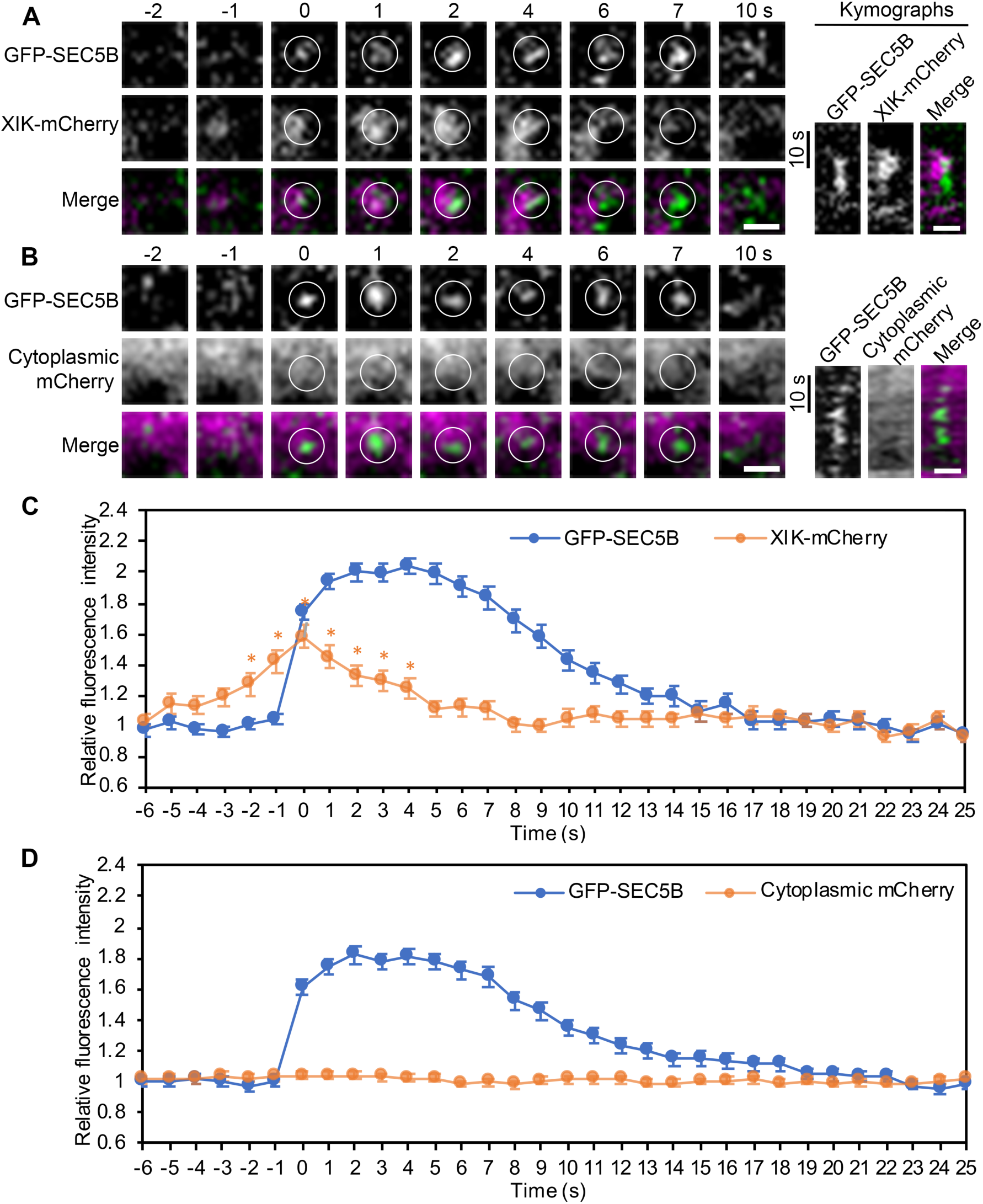
Myosin XIK Transiently Colocalizes with Stationary Foci of SEC5B at the PM. **(A)** Representative time series show arrival of a GFP-SEC5B particle (white circles, green in merged images) at the PM that was colocalized with XIK-mCherry (magenta) in an etiolated hypocotyl epidermal cell imaged with spinning disk confocal microscopy. The images and corresponding kymographs show that colocalization occurred transiently in the first few seconds upon arrival of the GFP-SEC5B particle. Bars = 1 µm. **(B)** Representative time series show localization of a GFP-SEC5B particle (white circles, green in merged images) and cytoplasmic mCherry (magenta) signal near the PM. The images and corresponding kymographs show that the mCherry signal was constantly present at low levels throughout the entire time course without showing any specific association with the GFP-SEC5B particle. Bars = 1 µm. (**C, D**) Quantitative analysis of fluorescence intensity for newly-appearing stationary foci of GFP-SEC5B at the PM in the GFP channel and the corresponding XIK-mCherry (**C**) or cytoplasmic mCherry signal (**D**) in the mCherry channel. There was significantly higher fluorescence intensity of XIK-mCherry that peaked at 0 s and lasted for 4 s after the appearance of new GFP-SEC5B particles. In contrast, there were no significant changes in fluorescence intensity of cytoplasmic mCherry that correspond to newly arrived GFP-SEC5B particles. Values given are means ± SE (For intensity assay in **C**, n = 71 particles; for intensity assay in **D**, n = 78 particles; the X-bar and S Control Charts were used for statistical comparison of timepoints; *P < 0.05).

In addition to colocalization of XIK with SEC5B, we performed colocalization analysis of XIK with a second exocyst subunit, EXO70A1. We created a double marked line expressing XIK-mCherry and EXO70A1-GFP and observed a similar spatiotemporal pattern of colocalization which only occurred in the first few seconds upon arrival of the EXO70A1 foci (Supplemental Figure 4).

Considering that XIK-mCherry mostly appeared as clusters of diffuse, amorphous signal in the cortical cytoplasm, and to exclude the possibility that the colocalization with exocyst was due to non-specific cytoplasmic signal, we co-expressed a cytoplasmic mCherry construct with GFP-SEC5B and conducted colocalization analysis. We observed that in most cases, a constant mCherry signal was present at the same region with the newly-appeared SEC5B particle throughout the time course (Figure 5B). Fluorescence intensity analysis showed that there was no significant difference of fluorescence intensity of cytoplasmic mCherry among all the time points measured in the selected ROIs that correspond to the SEC5B foci (Figure 5D). Thus, these results indicate that there was specific association of XIK with the exocyst complex near the PM, presumably at an early stage of vesicle tethering.

### Exocyst Localization and Lifetime at the Vesicle Tethering Site in CSC Secretion Depend on Myosin XIK

To further confirm the functional relationship between XIK and the exocyst complex in vesicle tethering, we investigated their association during exocytosis of CSCs at the PM. Previous studies show that both SEC5B (Zhu et al., 2018) and XIK (Zhang et al., 2019) transiently associate with CSCs at the beginning of the pause phase during CSC secretion; mutation of either exocyst subunits or XIK resulted in reduced CSC insertion/delivery rates or increased insertion defects, suggesting both components are critical for CSC secretion at the PM. Here, we first verified the spatiotemporal association of exocyst and XIK with CSCs during the vesicle tethering step. We tested two exocyst subunits, SEC5B and EXO70A1, using double marked lines of GFP-SEC5B tdTomato-CESA6 *prc1-1* and EXO70A1-GFP tdTomato-CESA6 *prc1-1*, as well as the colocalization of XIK with CESA6 using the double marked line of XIK-mCherry YFP-CESA6 *prc1-1 xik-2.* We tracked single CSC insertion events during the progression from erratic phase, pause phase, to the steady translocation phase in the double marked lines and measured the fluorescence intensity in the corresponding exocyst or XIK channels (Figure 6; Supplemental Figure 5).

**Figure 6.**
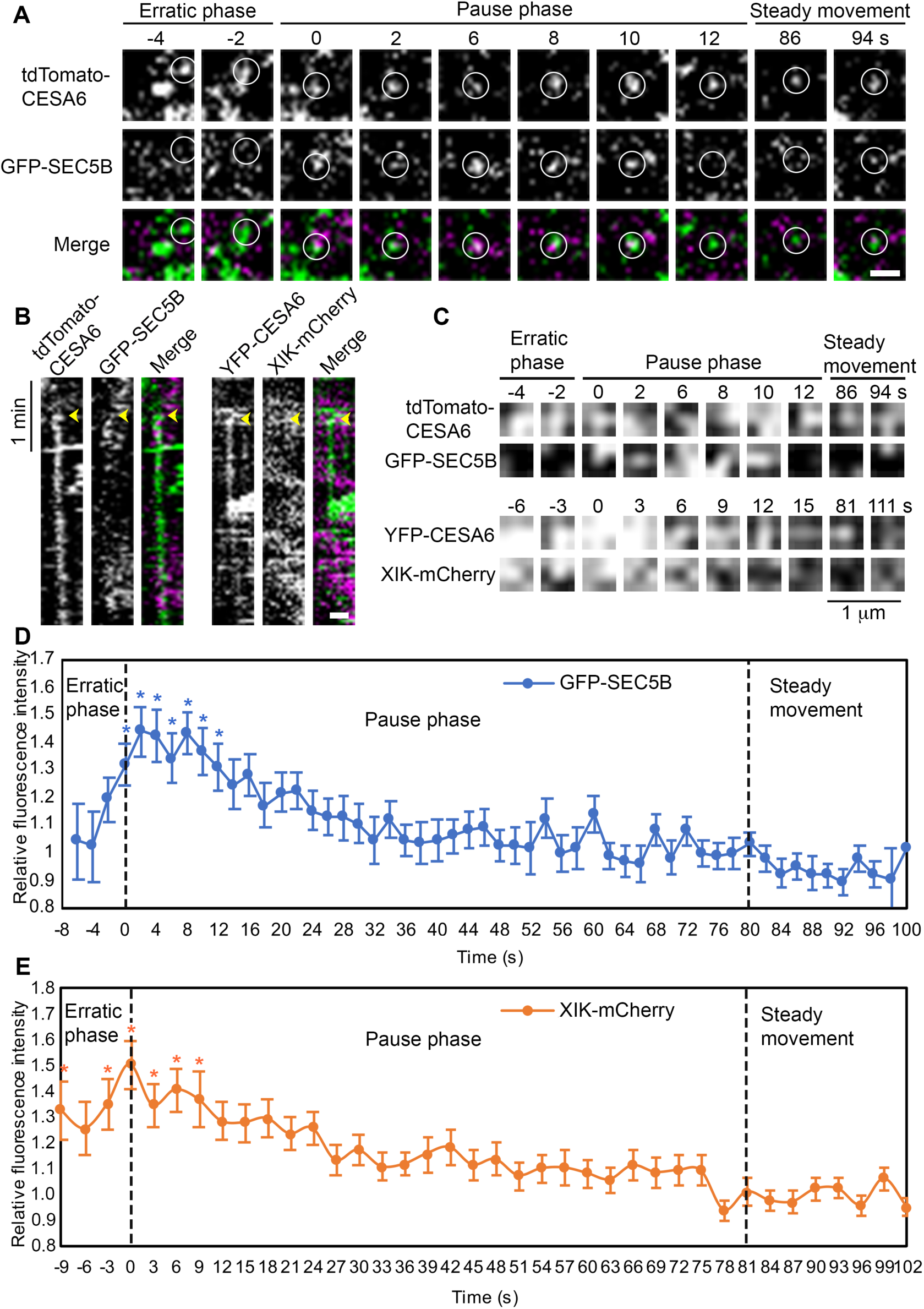
Myosin XIK and Exocyst Subunits Colocalize with CESA6 during the Vesicle Tethering Step of Secretion. **(A)** Representative images of a time series collected from the cortical cytoplasm of a hypocotyl epidermal cell show that GFP-SEC5B (magenta) colocalized with a newly-arrived CSC vesicle (white circles, green in merged images) during the first few seconds of the pause phase, but not during the erratic phase or steady movement phase. Bar = 1 µm. **(B)** Representative kymographs demonstrate transient colocalization of GFP-SEC5B (magenta) with tdTomato-CESA6 (green), as well as XIK-mCherry (magenta) with YFP-CESA6 (green) at the beginning of the pause phase (yellow arrowheads). Bar = 1 µm. **(C)** A small ROI (3 × 3 pixels) was selected at the centroid of a single CSC vesicle or particle from the representative images shown in **(A)**. The ROI, defined by the presence of CESA6, was tracked in both channels during the erratic phase, pause phase, and the steady movement phase. The ROI in the GFP-SEC5B channel or the XIK-mCherry channel was extracted for analysis of fluorescence intensity. **(D, E)** Quantitative analysis of mean fluorescence intensity of GFP-SEC5B (**D**) or XIK-mCherry (**E**) in ROIs as shown in (**C**) from multiple insertion events in different epidermal cells. The fluorescence intensity was normalized using the intensity of the ROI in each frame divided by the average intensity of ROIs from the steady movement phase, assuming that any signal associated with an actively translocating CSC represents a random event. Values given are means ± SE (For GFP-SEC5B intensity assay, n = 40 insertion events, because the duration of the erratic phase varied among different insertion events, the sample sizes at −6 s, −4 s, and −2 s were 12, 10, and 21, respectively; for XIK-mCherry intensity assay, n = 63 insertion events, and the sample sizes at −9 s, −6 s, and −3 s were 19, 31, and 49, respectively; the X-bar and S Control Charts were used for statistical comparison of timepoints; *P < 0.05).

Similar to the previous report (Zhu et al., 2018), we observed that in most insertion events, a GFP-SEC5B particle appeared at the location of a CSC particle at the PM at the beginning of the pause phase (Figures 6A to 6C; Supplemental Movie 5), and the colocalization lasted for an average duration of 11.2 ± 5 s (n = 115 insertion events). The colocalization was further confirmed by fluorescence intensity analysis over a time course in the GFP-SEC5B channel in small ROIs that correspond to the centroid of a CSC particle (Figures 6C and 6D). Significantly higher fluorescence signal was detected from 0 to 12 s after the beginning of the pause phase compared with the remaining time points (Figure 6D). The colocalization of EXO70A1-GFP with tdTomato-CESA6 exhibited a similar spatiotemporal pattern, with the association of EXO70A1-GFP signal at the beginning of the pause phase for 12.8 ± 6 s (n = 90 insertion events; Supplemental Figure 5). We next examined the colocalization of XIK-mCherry with YFP-CESA6 during CSC secretion. Similar to the earlier study (Zhang et al., 2019), we detected transient colocalization of XIK-mCherry with YFP-CESA6 during the erratic phase, when a CSC vesicle initially arrives near the PM, as well as during the first few seconds at the beginning of the pause phase (Figures 6B and 6C). Due to the diffuse pattern of the XIK-mCherry signal in the cell cortex as described above, the specific association of XIK with CSC particles was confirmed by fluorescence intensity quantification in the XIK-mCherry channel. There was significantly higher signal of XIK-mCherry during the erratic phase and the first 9 s at the beginning of the pause phase (Figure 6E), suggesting XIK was specifically associated with CSC particles at those time points.

In contrast to the observation that XIK was associated with the erratically moving vesicles in the cortex before the pause and insertion of CSCs, we failed to detect any significant colocalization of either GFP-SEC5B or EXO70A1-GFP foci with CSCs during the erratic phase (Figures 6A to 6D; Supplemental Figure 5; Supplemental Movie 5). In yeast, some models suggest that EXO70 and SEC3 directly bind to the PM and other subunits arrive at the PM subsequently by interaction with secretory vesicles (Boyd et al., 2004; Wu and Guo, 2015). In plants, the dynamics of exocyst complex formation during vesicle secretion remain largely unknown (Saeed et al., 2019). Our results suggest that at least the SEC5B or EXO70A1 subunits were not pre-attached to secretory vesicles before PM tethering and fusion, rather, both subunits arrived at the CSC insertion site from the nearby cortex or membrane after the arrival of a CSC and myosin XIK, which is different from the yeast model. Another possibility is that the temporal resolution (2-s intervals) in this study was unable to detect some more transient colocalization events or sequential arrival events (< 2 s). Nevertheless, our results show that during a CSC insertion event, XIK associated with a CESA compartment from the erratic phase to the initial tethering phase, whereas the exocyst appears to arrive at a later time, only from the beginning of the tethering phase.

Because XIK associates with CESA compartments in advance of exocyst subunits during exocytosis and because disruption of myosin XI activity reduces the frequency and lifetime of stable exocyst foci at the PM, we tested whether myosin activity is required for the localization and dynamics of exocyst at vesicle tethering sites during CSC delivery. We generated double marked lines expressing GFP-SEC5B tdTomato-CESA6 and EXO70A1-GFP tdTomato-CESA6 in the *xik-2* homozygous background as well as wild-type siblings expressing both reporters. We also applied pre-treatment with PBP for 10 min to acutely inhibit myosin XI activity in wild-type seedlings. The CSC insertion events were tracked in double marked lines and only successful insertion events were quantified, assuming those events rely on the exocyst complex during the vesicle tethering step. The colocalization of exocyst subunits with CSC particles was examined during tethering and only exocyst foci that were continuously present for 2 frames (4 s) or longer at the insertion site from the beginning of the pause phase were considered colocalized. With the GFP-SEC5B tdTomato-CESA6 co-expression line, we detected positive association of SEC5B foci with CESA6 at the beginning of the pause phase in an average of 87% of the insertion events from 3 independent experiments, however, the average colocalization rate dropped significantly to 71% and 59% in *xik-2* and PBP-treated cells, respectively (Figures 7A and 7B). The reduced rate of colocalization was further confirmed by fluorescence intensity analysis. In wild type, the average SEC5B fluorescence intensity was significantly higher from 0 to 12 s at the beginning of the pause phase compared with the rest of the time points (Figure 7C), which was consistent with the average lifetime of exocyst measured in our earlier results and indicated a high frequency of association of SEC5B with CESA6 during the first 12 s of the pause phase. In contrast, decreased fluorescence intensity was detected in *xik-2* and PBP-treated cells during the first 12 s of the pause phase (Figure 7C). In *xik-2*, only the 0, 2, and 6 s time points showed significantly higher fluorescence signal, and in PBP-treated cells, none of the time points tested were significantly different compared with the other time points (Figure 7C), suggesting that there was no significant colocalization of SEC5B foci with CESA6 at the tethering phase in those cells.

**Figure 7.**
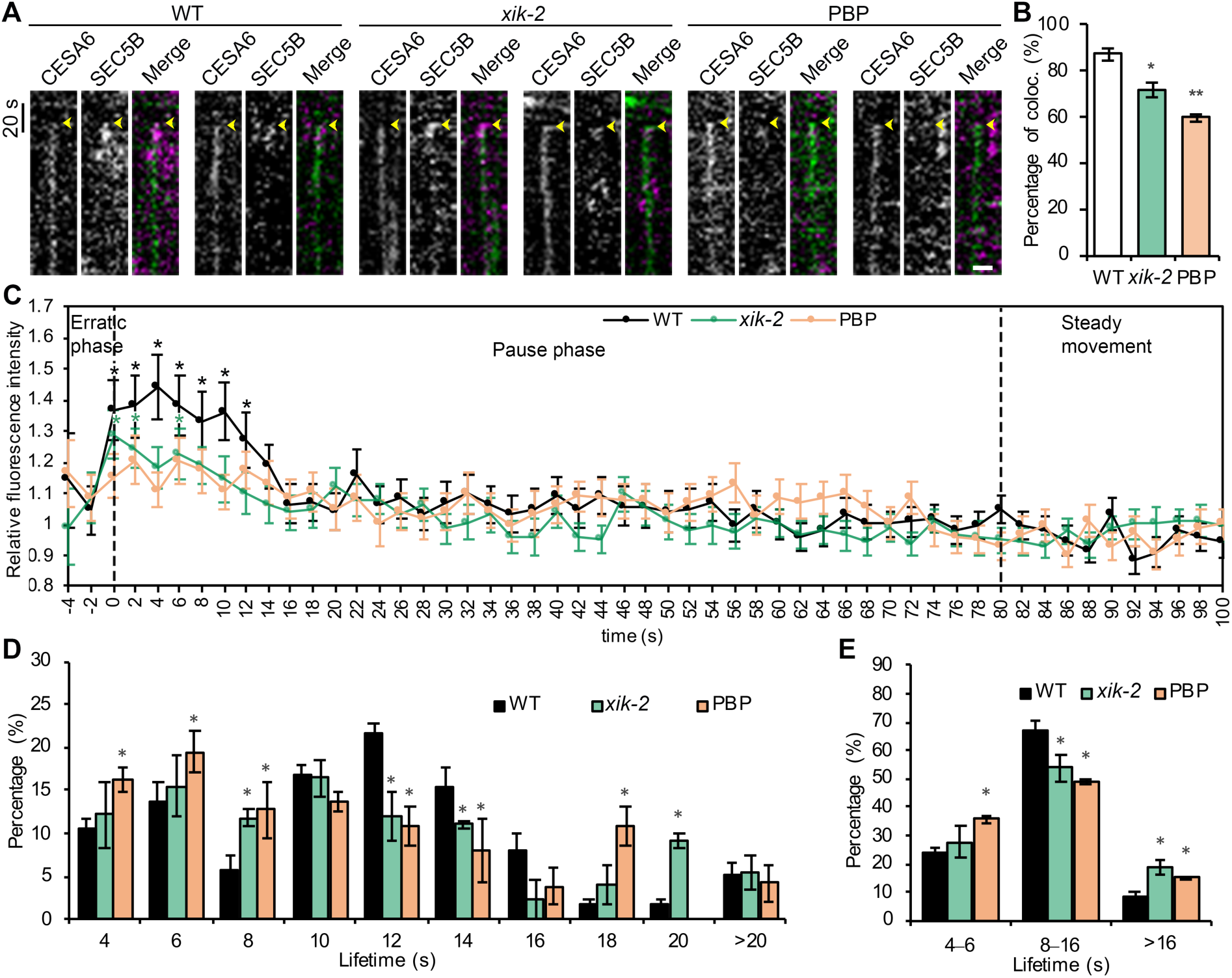
Disruption of Myosin Activity Results in Reduced Colocalization of SEC5B with CESA6 and Altered SEC5B Tethering Time at the PM during CSC Secretion. **(A)** Representative kymographs show colocalization of GFP-SEC5B (magenta) and tdTomato-CESA6 (green) at the beginning of the pause phase (yellow arrowheads) during CSC insertion events. Cells were treated with mock (0.5% DMSO) or 10 μm PBP for 10 min prior to dual-channel time-lapse imaging. In *xik-2* or PBP-treated cells, the lifetime of GFP-SEC5B appeared shorter or there was no SEC5B foci colocalized with the CESA6 particle at the pause phase. Bar = 1 µm. **(B)** Quantitative analysis shows that the percentage of colocalization between SEC5B and CESA6 at the beginning of the pause phase was greatly reduced in *xik-2* and PBP-treated cells. Values given are means ± SE (n = 3 biological repeats; A total of 154, 130, and 134 CSC insertion events were tracked in WT, *xik-2* and PBP-treated cells, respectively; Student’s t test, *P < 0.05, **P < 0.01). **(C)** Quantitative analysis of mean fluorescence intensity of GFP-SEC5B in ROIs that correspond to the centroid of a CSC particle during the erratic phase, pause phase, and the steady movement phase. Values given are means ± SE (Data represents one biological repeat; n = 44, 43, and 37 insertion events in WT, *xik-2* and PBP-treated cells, respectively; the X-bar and S Control Charts were used for statistical comparison of timepoints; *P < 0.05). (**D**, **E**) Distribution of lifetimes of GFP-SEC5B foci at the vesicle tethering sites during secretion. The distribution of lifetimes was further grouped into three subpopulations (**E**) based on overall mean ± SD (11.5 ± 5 s) measured in wild type with the same data shown in (**D**). Values given are means ± SE (n = 3 biological repeats; A total of 115, 99, and 73 SEC5B foci were measured in WT, *xik-2* and PBP-treated cells, respectively; Student’s t test, *P < 0.05).

The lifetime of GFP-SEC5B at CSC insertion sites was also reduced in *xik-2* or PBP-treated cells, compared with that in wild-type cells (Figures 7A, 7D, and 7E). In wild type, the distribution of the resident lifetimes of GFP-SEC5B revealed a non-Gaussian distribution and three subpopulations: in a majority of colocalization events (∼70%), a GFP-SEC5B foci was associated with CESA6 for 8–16 s (defined based on overall mean ± SD of 11.5 ± 5 s for wild type); ∼20% of the events only lasted for 4–6 s, and the remaining 10% showed colocalization for 18 s or more (Figures 7D and 7E). However, in *xik-2* or PBP-treated cells, the proportion of colocalization events that had a lifetime of 8–16 s was decreased by 15–20%, whereas the populations with shorter (< 8 s) or prolonged (>16 s) duration of SEC5B association were increased in myosin-deficient cells, compared with that in wild type (Figures 7D and 7E).

Because yeast EXO70 is proposed to interact with the PM to mark secretion sites and recruits other subunits together with the secretory vesicle to the destination membrane (Boyd et al., 2004; Wu and Guo, 2015), we examined whether the localization and dynamics of EXO70A1 was affected by myosin XI activity during CSC secretion. Using the EXO70A1-GFP tdTomato-CESA6 co-expression line, we detected a reduction of the colocalization rate of EXO70A1-GFP with CESA6 at the tethering stage in both *xik-2* and PBP-treated cells, comparable to that observed for SEC5B in the previous experiment (Figure 8). The lifetime analysis of EXO70A1-GFP foci at the CSC insertion sites showed that there was a significantly increased proportion of insertion events that only had a transient exocyst association of 4 s and the lifetime peaked at 8–10 s in *xik-2* and PBP-treated cells, compared with that in wild-type cells which had a peak colocalization duration for 12–14 s (Figure 8). The results suggest that similar to SEC5B, the association and lifetime of EXO70A1 during vesicle tethering is also dependent on myosin XI.

**Figure 8.**
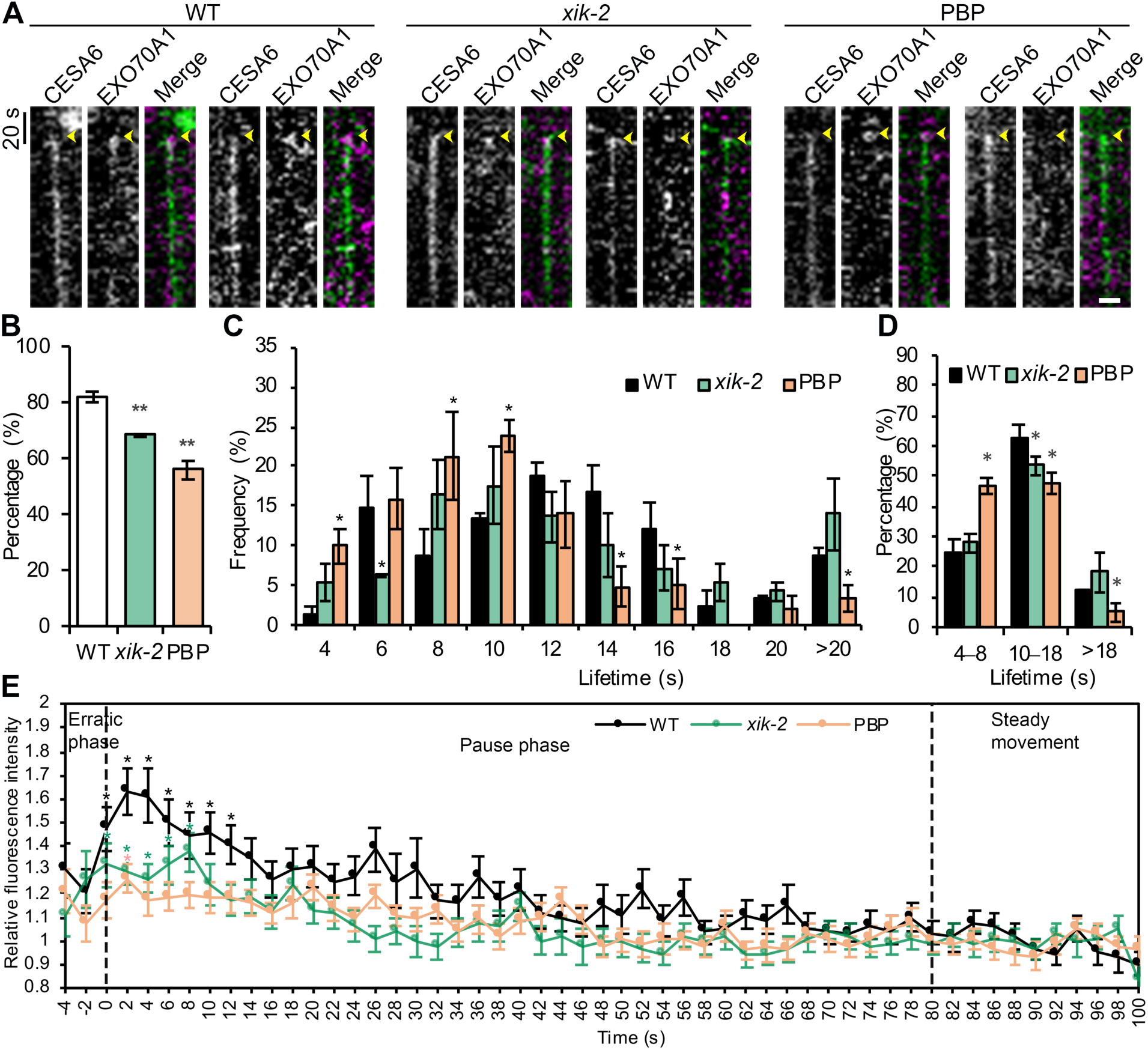
Disruption of Myosin Activity Results in Reduced Colocalization of EXO70A1 with CESA6 and a Shorter EXO70A1 Tethering Time during CSC Secretion. **(A)** Representative kymographs show colocalization of EXO70A1-GFP (magenta) and tdTomato-CESA6 (green) at the beginning of the pause phase (yellow arrowheads) during CSC insertion events. Cells were treated with mock (0.5% DMSO) or 10 μm PBP for 10 min prior to dual-channel time-lapse imaging. In *xik-2* or PBP-treated cells, the lifetime of EXO70A1-GFP appeared shorter or there was no EXO70A1 foci colocalized with the CESA6 particle at the pause phase. Bar = 1 µm. **(B)** Quantitative analysis shows that the percentage of colocalization between EXO70A1 and CESA6 at the beginning of the pause phase was significantly reduced in *xik-2* and PBP-treated cells. Values given are means ± SE (n = 3 biological repeats; A total of 141, 162, and 96 CSC insertion events were tracked in WT, *xik-2* and PBP-treated cells, respectively; Student’s t test, **P < 0.01). (**C, D**) Distribution of lifetimes of EXO70A1-GFP foci at the vesicle tethering sites during secretion. The distribution of lifetimes was further grouped into three subpopulations (**D**) with the same data shown in (**C**). Values given are means ± SE (n = 3 biological repeats; A total of 120, 97, and 69 EXO70A1 foci were measured in WT, *xik-2* and PBP-treated cells, respectively; Student’s t test, *P < 0.05). (**E**) Quantitative analysis of mean fluorescence intensity of EXO70A1-GFP in ROIs that correspond to the centroid of a CSC particle during the erratic phase, pause phase, and the steady movement phase. Values given are means ± SE (Data represents one biological repeat; n = 45, 35, and 23 insertion events in WT, *xik-2* and PBP-treated cells, respectively; the X-bar and S Control Charts were used for statistical comparison of timepoints; *P < 0.05).

Collectively, our data demonstrate that disruption of myosin XI inhibited the localization and lifetime of exocyst subunits at vesicle tethering sites during the exocytosis of CSCs. Using CESA as a model cargo, our results confirmed that plant myosins and exocyst complex cooperate to mediate vesicle tethering near the PM and the localization and dynamic behavior of exocyst at the site of exocytosis is dependent upon myosin XI activity, as shown in Figure 9.

**Figure 9.**
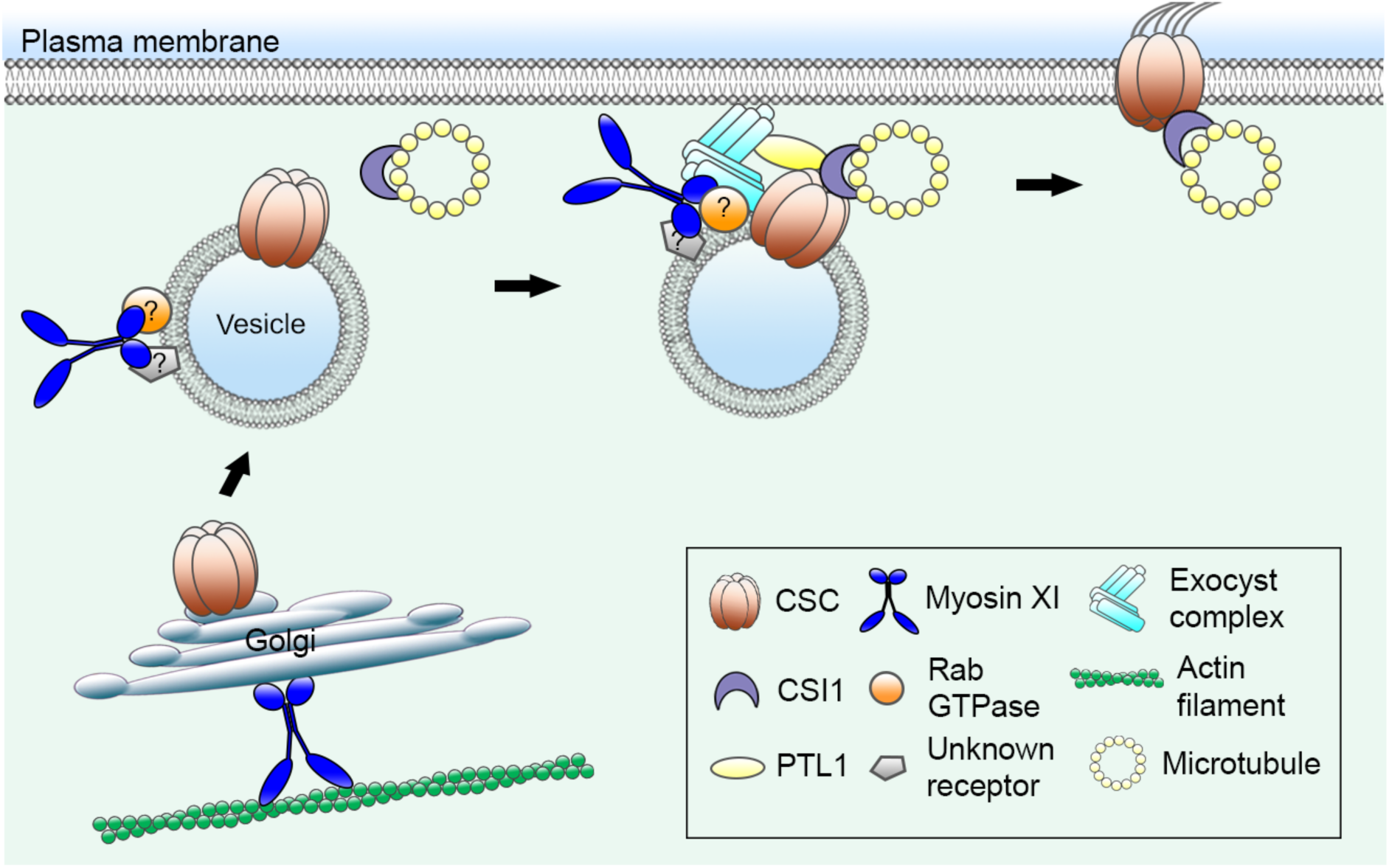
Roles for Myosin XI in Delivery of CSCs to the PM. Myosin XI mediates CSC delivery at two levels: One group of motors is responsible for the cell-wide transport and distribution of CESA-containing Golgi; another group is transiently associated with secretory vesicles potentially through interactions with an unknown receptor and a Rab GTPase. Once a secretory vesicle is anchored along cortical microtubules through an interaction between CESA and CSI1, the myosin XI and possibly a Rab GTPase on the vesicle surface are required for the recruitment and stabilization of exocyst complex subunits at the PM site for membrane tethering and fusion. Vesicle exocytosis is also facilitated by interactions between CSI1, PTL1, and exocyst subunits, as shown previously (Zhu et al., 2018).

## DISCUSSION

Myosin XI motors power movement along the actin cytoskeleton and are major contributors to the transport and distribution of intracellular components in plant cells, thereby playing a key role in regulating cell growth and development. In a previous study, we demonstrated a new role for myosin XI in exocytosis through regulating vesicle tethering or fusion, although the exact mechanism was unclear (Zhang et al., 2019). Here, we showed that myosin XIK, the predominant motor driving organelle transport in plant vegetative cells, participates in the vesicle tethering step of exocytosis through direct interactions with the exocyst complex via its globular tail domain (GTD). Specifically, myosin XIK GTD bound directly to SEC5B in vitro and a functional fluorescently-tagged XIK colocalized with multiple exocyst subunits at PM-associated stationary foci. Moreover, genetic and pharmacological inhibition of myosin activity reduced the frequency and lifetime of stationary exocyst complexes, which are presumptive sites of vesicle tethering and docking during secretion. Using high spatiotemporal resolution imaging and pair-wise colocalization analysis of myosin XIK, exocyst subunits, and CESA6 in single CSC exocytosis events, we demonstrated that XIK associates with secretory vesicles earlier than exocyst and likely recruits the exocyst to the PM tethering site to initiate vesicle tethering. This study provides new insights about the dynamic regulation of exocytosis in flowering plants as well as the role of plant myosin XI in secretion.

### A Conserved Role for Myosin XI in Exocytosis

Our results reveal an evolutionarily-conserved role for myosin XI that includes direct interaction with exocyst complex subunits and participation in exocytosis, similar to the well characterized yeast myosin V motor, Myo2p. In budding yeast, Myo2p delivers secretory vesicles to the growing bud and binds directly to the SEC15 subunit of exocyst through conserved amino acids at its tail cargo-binding domain (Jin et al., 2011). The Rab GTPase SEC4 binds to Myo2p as well as SEC15 and is thought to recruit SEC15 and thereby other exocyst subunits to the secretory vesicle surface (Guo et al., 1999; Jin et al., 2011; Santiago-Tirado et al., 2011). Mutation of SEC15-binding sites on Myo2p results in failed localization of SEC15 to the growing bud tip, suggesting that Rab activity alone is not sufficient for recruiting exocyst to the vesicles and Myo2p is also required for the correct localization of exocyst during exocytosis. Consistent with the yeast model, we showed that during secretion of CSCs, myosin XIK was required for the localization of SEC5B and EXO70A1 to CSC insertion sites. Combined with results that an overall reduction of membrane dwelling events of SEC5B, EXO70A1 and SEC6 were observed in *xik* or PBP-treated cells, and evidence that XIK colocalized with newly-appeared exocyst subunits at the PM, we propose that XIK may have a general role in recruiting exocyst complex to exocytosis sites, not just for CSC trafficking.

Homology modeling reveals a C-terminal cargo binding domain structure with similarities between Arabidopsis myosin XI and Myo2p (Li and Nebenführ, 2007). The plant exocyst complex also consists of 8 conserved subunits that have similar rod-like structures as shown for yeast and mammalian subunits (Elias et al., 2003; Hála et al., 2008; Žárský et al., 2013). Therefore, it is not surprising that myosin– exocyst interactions also occur in plant cells. However, the plant proteins may have evolved divergent interactions or functions. In contrast to Myo2p, we showed that myosin XIK GTD did not interact with SEC15 in Arabidopsis, but interacted strongly with SEC5B and exhibited a weaker interaction with Exo84A in a yeast two-hybrid screen. The interaction with SEC5B was confirmed with an in vitro pull-down assay. In yeast, SEC15 is considered one of the most proximal subunits to secretory vesicles as it directly interacts with the Rab SEC4, which also binds to Myo2p at the vesicle surface (Guo et al., 1999; Jin et al., 2011). Therefore, this SEC4-Myo2p-SEC15 protein complex brings together secretory vesicles, motors and the exocyst complex to couple polarized vesicle transport to exocytosis. In plants, it is still unknown which exocyst subunit is responsible for secretory vesicle binding and which Rab regulates exocyst dynamics during exocytosis. Interestingly, Arabidopsis SEC5B was also shown to interact with CESA6 in a yeast two-hybrid assay (Zhu et al., 2018), together with our results that SEC5B interacts with XIK, it is highly likely that in plant cells, SEC5B represents one of the subunits that mediates secretory vesicle binding. A recent study shows that Arabidopsis SEC15B interacts with STOMATAL CYTOKINESIS DEFECTIVE1 (SCD1) and SCD2, which also interact with RabE1, a close homolog of SEC4 in plants, suggesting a potential interaction that brings together Rabs and the exocyst complex in post-Golgi trafficking (Mayers et al., 2017). In plant cells, myosin XI, exocyst and Rab gene families are all expanded in size compared to that in the yeast, and it is plausible that plants utilize a more complicated and specific interaction network during exocytosis to fulfill the needs of different cell types and trafficking pathways. Further studies are needed to uncover the functional connection between myosin motors, exocyst, Rab GTPases and other players in the highly regulated secretory trafficking processes in plants.

### Dynamics of the Exocyst Complex During Vesicle Tethering

Although recent studies show that the exocyst complex plays important roles in many secretion-related processes in plants, such as root hair and hypocotyl elongation, cell division, cell wall deposition, auxin signaling and defense response against pathogens (Synek et al., 2006; Hála et al., 2008; Fendrych et al., 2010; Drdová et al., 2013; Žárský et al., 2013; Vukašinović et al., 2017; Pečenková et al., 2020), the assembly and dynamic regulation of the exocyst complex during vesicle tethering remain enigmatic. Different models for the dynamic assembly of exocyst during exocytosis have been proposed in yeast and mammalian cells. In budding yeast, the EXO70 and SEC3 subunits are localized at the PM through binding with membrane lipids (Boyd et al., 2004; He et al., 2007; Pleskot et al., 2015), whereas other subunits are vesicle-bound and when the vesicle arrives at the secretion sites, the PM-bound subunits interact with the vesicle-bound population to form a holocomplex that tethers the vesicle to the PM (Donovan and Bretscher, 2015; Mei and Guo, 2019). However, plant cells seem to have distinct mechanisms for exocyst assembly during vesicle tethering. Using CESA6 as a vesicle marker, we captured single vesicle tethering events with SDCM and tracked the dynamics of two exocyst subunits, SEC5B and EXO70A1. Both SEC5B and EXO70A1 appeared at the CSC insertion site coincident with the stabilization of CSC vesicles at the PM and had a similar average lifetime of 11–12 s; disruption of myosin XI equally affected the localization of the two subunits to the PM insertion sites. Further, neither subunit was pre-associated with the vesicles prior to tethering and docking, whereas myosin XIK arrived with the CSC vesicle and showed erratic movement with the compartment before it became stationary. Myosin XIK also associated transiently with the tethering site and had a lifetime of 3–9 s. These results suggest that EXO70A1 does not pre-exist at the PM secretion site to recruit other vesicle-bound subunits as suggested in the yeast model, and neither SEC5B and EXO70A1 are delivered to the PM on secretory vesicles. Consistent with our findings, a previous study that tracked several exocyst subunits with high resolution VAEM shows that exocyst subunits reside at the PM as dense particles whose density is obviously higher than the expected number of vesicle tethering/exocytosis events, and their localization at the PM is independent of secretory vesicle or exocytosis, as brefeldin A treatment which blocked secretion did not affect the density of exocyst foci at the PM (Fendrych et al., 2013). In addition, exocyst subunit density at the PM is also unaffected by short-term perturbation of cortical actin and microtubules by inhibitor treatment (Fendrych et al., 2013). Another study shows that the PM density of Sec3A is similar among growing and non-growing cells in interphase cells and Sec3A subunits do not preferentially accumulate in PM regions that undergo bulk exocytosis (Zhang et al., 2013). Similarly, we showed that the overall density and distribution pattern of SEC5B, EXO70A1 and SEC6 foci at the PM were unchanged in *xik* or cells with short-term PBP or LatB treatment, even though those cells were supposed to have a reduction in overall exocytosis rate (Zhang et al., 2019). Based on these data, it is highly likely that in plants, the EXO70 and SEC3 subunits do not function as landmark proteins to recruit other subunits during vesicle tethering, instead, most plant exocyst subunits pre-exist at the PM regardless of the presence of vesicles, and are transiently recruited to the secretion sites when vesicles arrive at the PM.

Fendrych et al. (2013) also propose that the exocyst foci at the PM represent pre-assembled complexes, although direct evidence is lacking. Several recent studies in yeast and mammalian cells suggest that exocyst can assemble into two stable subcomplexes (SEC3-SEC5-SEC6-SEC8 and SEC10-SEC15-EXO70-EXO84) and a new model shows that upon arrival of secretory vesicles to the PM, the two subcomplexes are triggered to assemble into a holocomplex to tether the vesicle to the membrane (Heider et al., 2016; Ahmed et al., 2018; Mei et al., 2018; Mei and Guo, 2019). In plants, it remains to be determined whether the PM-localized exocyst subunits are pre-assembled into similar subcomplexes or a holocomplex. Nevertheless, it is plausible that in plants, the exocyst subunits reside at the PM either as pre-assembled complexes, or are assembled immediately upon arrival of the secretory vesicle at the PM sites, and in either case, the subunits are recruited to the secretion sites by myosin motors and possibly an unknown Rab GTPase on the vesicle surface. Further, we show that myosin XI affects the lifetime of exocyst at the PM tethering site during CSC insertion. Our results provide the first detailed study of exocyst tethering time in verified exocytosis events in plant cells, as documented by the insertion of a functional cellulose-synthesizing complex into the PM. We observed, based on the distribution of lifetimes of both SEC5B and EXO70A1 during CSC insertion, that there are three types of association of exocyst during exocytosis. The majority have an association for 10–16 s, suggesting those likely represent standard tethering events. This duration is very similar to the reported exocyst residency time during exocytosis in yeast and mammalian cells of 12–18 s (Donovan and Bretscher, 2015; Ahmed et al., 2018), indicating a conserved vesicle tethering/fusion process in all eukaryotes. A short exocyst association of 4–6 s was also observed in ∼20% of the events, likely representing a short-lived or aborted exocyst complex that failed to form a mature complex, similar to that reported for the formation of clathrin-coated pits for endocytosis (Loerke et al., 2009). Notably, we found that this short-lived population was significantly increased in cells with compromised myosin XI activity. Similarly, the average lifetime of several subunits was reduced by 1–2 s in myosin-deficient cells in general. Although the detailed mechanism is unclear, one possibility is that myosin XI is required for the formation of stable exocyst complexes at the PM. A third population of exocyst (∼10%) has a lifetime of 20 s or longer at CSC tethering sites, however, it is unclear whether those represent prolonged or defective vesicle tethering or fusion events. The regulation of exocyst dynamics during vesicle tethering remain somewhat unclear and require further investigation with additional markers and higher spatiotemporal resolution imaging, as accomplished in mammalian cells (Ahmed et al., 2018, Mei and Guo, 2019). Besides myosin XI, other protein components have been shown to interact with exocyst subunits and may coordinate exocyst dynamic function during secretion. CESA6 interacts with SEC5B in a yeast two-hybrid assay, indicating that cargo proteins on the secretory vesicle surface may also play a role in recruiting or stabilizing the exocyst complex at the PM–vesicle interface (Zhu et al., 2018). A plant specific protein PATROL1 (PTL1) directly interacts with SEC10 and is required for the exocytosis of CSCs (Zhu et al., 2018). PTL1 arrives at the CSC insertion sites 1– 2 s later than SEC5B and genetic mutation of PTL1 did not affect SEC5B dynamics, suggesting that PTL1 likely has a role in a later step during exocytosis. Another protein with a probable role during the fusion step of exocytosis, the Sec1/Munc18-related protein KEULE, has been shown to interact with SEC6 during cell plate formation (Wu et al., 2013). Finally, similar to the situation in yeast, interactions between SNAREs and exocyst subunits have been demonstrated in plants (Larson et al., 2020). Collectively, these data suggest that several multiprotein complexes must cooperate in spatiotemporal manner during the CSC pause phase to build the macromolecular machines that execute vesicle tethering, docking, and fusion.

### A New Model for Post-Golgi Trafficking Regulated by the Cytoskeleton

While the actin–myosin XI transport network plays a predominant role in long-distance organelle and vesicle movement, its role in local post-Golgi or secretory vesicle trafficking is less clear (Nebenführ and Dixit, 2018). In tip-growing cells (root hairs, pollen tubes and polarized moss cells), actin and myosin XI are proposed to directly regulate the targeted delivery of secretory vesicles to the growing apex (Park and Nebenführ, 2013; Madison et al., 2015; Orr et al., 2020), whereas in diffusely-growing cells, a similar role has not been established. Our results from diffusely-growing Arabidopsis epidermal cells demonstrate that myosin XI and actin play a direct and active role in the final steps of secretory vesicle trafficking by recruiting the exocyst complex and mediating vesicle tethering at the PM, rather than during an early vesicle transport step (Figure 9).

Although myosin XIK was shown to be responsible for the motility of several compartments potentially involved in secretion, such as the trans-Golgi network, putative secretory vesicles, and endosomes in diffuse growing cells (Avisar et al., 2012; Peremyslov et al., 2015), it is not known whether myosin binds directly to these compartments or mediates the targeted delivery of these compartments to specific PM regions for secretion. Instead, indirect or passive models are favored by some; specifically, it is proposed that vesicles and organelles move passively with the hydrodynamic flow generated by myosin XI-driven cytoplasmic streaming (Peremyslov et al., 2013; Buchnik et al., 2015; Nebenführ and Dixit, 2018). Evidence supporting these indirect models came from the study of a major group of plant-specific myosin receptors, MyoBs, which attach myosin XI to a specific type of small endomembrane compartment to drive rapid cytoplasmic streaming, and these structures do not colocalize with any known organelle or vesicle markers (Peremyslov et al., 2013; Peremyslov et al., 2015). Early studies expressing dominant-negative myosin XI tail constructs also showed no major colocalization with organelles or vesicles, although their motility was inhibited (Sparkes et al., 2008; Avisar et al., 2009; Avisar et al., 2012).

Specific secretory vesicle receptors such as Rab GTPases that directly link myosin XI to cargo and support a role for myosin XI in secretory vesicle transport have not been identified in flowering plants. In budding yeast, the Rab GTPase SEC4 plays a central role in polarized transport of secretory vesicles to the bud tip by interacting and recruiting myosin V motors and a range of other machinery proteins to the vesicle surface (Jin et al., 2011). One study in Arabidopsis identified two Rab GTPases, RabD1 and RabC2a, through a yeast two-hybrid screen using the myosin XI2 tail; however, RabC2a was shown to localize to peroxisomes and RabD mainly mediates ER to Golgi trafficking (Zheng et al., 2005; Hashimoto et al., 2008). A recent study identified RabE as a myosin XI partner in the moss *Physcomitrella patens* and this interaction is important for polarized growth (Orr et al., 2019). Further investigations are required to determine whether this interaction is conserved across different plant species and also functions in diffusely-growing cells. A Golgi localized RabH GTPase, RabH1b, has been shown to mediate the secretion of CSCs likely through regulating Golgi to PM trafficking, however, it is not known whether this process involves the interaction with cytoskeletal motors (He et al., 2018).

Finally, a wealth of evidence indicates a role for cortical microtubules and possibly kinesin motors, rather than actin and myosin, in mediating post-Golgi trafficking and membrane targeting of secretory vesicles (Nebenführ and Dixit, 2018; Elliott et al., 2020). Cortical microtubules and a kinesin-4 motor have been implicated in the trafficking of non-cellulosic cell wall components in Arabidopsis (Kong et al., 2015; Zhu et al., 2015). Cortical microtubules are also known to play an important role in CSC delivery by marking the CSC insertion sites through the linker protein CELLULOSE SYNTHASE INTERACTIVE1 (CSI1) and can interact with small CSC compartments that may be responsible for delivery or recycling of CSCs at the PM (Gutierrez et al., 2009; Bringmann et al., 2012; Li et al., 2012; Lei et al., 2015). CSI1 also interacts with PTL1, which is indicated to have a role in vesicle fusion in plants (Zhu et al., 2018). However, pharmacological removal of microtubules or genetic mutation of CSI1 showed no effect on CSC delivery rate to the PM (Gutierrez et al., 2009, Zhu et al., 2018, Zhang et al. 2019), indicating that CSI1 and microtubules may only serve as landmarks for the targeting of vesicles to the cortex and are not essential for subsequent steps like tethering and fusion with the PM. Further, in the *act2 act7* mutant as well as cells treated with actin or myosin inhibitors, a substantial reduction of CSC delivery/exocytosis rate was detected, however, the preferential positioning of CSCs to cortical microtubule sites was not affected, suggesting that the targeting of CSCs to cortical microtubules is independent of actin and myosin XI activity, precedes it in time and space, or both (Sampathkumar et al., 2013; Zhang et al., 2019).

In this study, we observed significantly higher signal of myosin XIK coincident with CESA compartments as early as 9 s before they arrive at PM insertion sites, even though it is unclear what receptors/adaptors mediate such an association or whether those motors play an active role in transport of CESA compartments to insertion sites that are marked by microtubules. Since the targeting of CSCs to cortical microtubules has been shown to be mediated by CSI1 (Zhu et al., 2018) and disruption of actin or myosin did not affect this targeting, myosin XIK on the vesicle surface may not have a major role in vesicle transport per se. It is likely that the force generated by cytoplasmic streaming is sufficient to propel the vesicles to the cortex or PM, or the CESA-containing Golgi transported along actin by myosin XI are already in close proximity to the PM to deliver newly formed secretory compartments. Indeed, previous models ascribe an indirect role for actomyosin in CSC delivery by maintaining a uniform global distribution of Golgi in the cortical cytoplasm of plant cells (Gutierrez et al., 2009; Sampathkumar et al., 2013). Identification of additional myosin XI receptors is required to confirm a role for myosin XI in active secretory cargo transport. Nevertheless, this study and recent work suggest that class XI myosins contribute to CSC delivery at two levels: one function is to power cytoplasmic streaming and cell-wide transport of Golgi bodies to the cortex or cortical microtubule sites, and once the delivery compartments are anchored to cortical microtubules, a second function of myosin XI is to cooperate with the exocyst complex and other players to facilitate local membrane tethering and fusion (Figure 9). This model is consistent with the finding that fluorescent-tagged functional myosin XIK exhibits two major locations in cells (Peremyslov et al., 2012; Zhang et al., 2019): one population displays a “beads-on-a-string” pattern to power cytoplasmic streaming and long-distance transport of organelles along actin cytoskeleton, whereas another population is more diffusely distributed in the cortical cytoplasm and is transiently associated with secretory vesicles to fulfill a conserved role in membrane tethering and exocytosis. Collectively, our study sheds new light on the spatiotemporal coordination of cytoskeleton and motors in regulating post-Golgi trafficking in plants, and helps uncover the evolutionally conserved and divergent regulation of exocytosis across kingdoms.

## METHODS

### Plant Materials and Growth Conditions

The Arabidopsis *myosin xi1 xi2 xik* triple knock-out (*xi3KO*) mutant and *xi3KO* expressing YFP-CESA6 in the homozygous *prc1-1* background were characterized previously (Peremyslov et al., 2010; Zhang et al., 2019). The *xi1*, *xi2*, and *xik-2* homozygous single mutant lines expressing YFP-CESA6 in the presence of *prc1-1* were recovered from the same cross that resulted in the previously characterized *xi3KO* YFP-CESA6 *prc1-1* lines (Zhang et al., 2019). The T-DNA insertion mutants *xik-1* (SALK_136682), *xik-2* (SALK_067972), *xi1* (SALK_019031) and *xi2* (SALK_055785) were obtained from the Arabidopsis Biological Resource Center (Ohio State University). Transgenic *Arabidopsis thaliana* Col-0 lines expressing EXO70A1-GFP and SEC6-GFP was described previously (Fendrych et al., 2010). To prepare GFP-SEC5B expressing lines, first, the Ubiquitin 10 promoter in the pUBN-GFP-DEST vector was restriction digested with *Sac*I and *Spe*I and replaced with the *SEC5B* native promoter (2000 bp upstream of the *SEC5B* start ATG) by using the NEB Gibson assembly master mix kit (MA, USA) to obtain a modified pSEC5BN-GFP-DEST vector. *SEC5B* promoter was amplified from Col-0 genomic DNA by primers TGACCATGATTACGAATTCGAGCTCTGTATTGAAACCCAAAATAT and CTCGCCCTTGCTCACCATACTAGTTGTTATCTCTGACTTAGATG. Then, the full-length CDS for *SEC5B* was cloned into the modified binary vector pSEC5BN-GFP-DEST using the Gateway system to obtain the final GFP-SEC5B expression vector. The XIK-mCherry construct was described previously (Peremyslov et al., 2012) and was kindly provided by Valerian V. Dolja (Oregon State University).

Double-marked lines were generated either by *Agrobacterium*-mediated transformation through floral dip or by crossing. The double-marked line for XIK-mCherry and EXO70A1-GFP was prepared by transforming the XIK-mCherry construct into plants expressing EXO70A1-GFP in the homozygous *xik-2* mutant background. For the XIK-mCherry and GFP-SEC5B double-marked line, XIK-mCherry in homozygous *xik-2* was crossed with GFP-SEC5B lines. For the double-marked line expressing GFP-SEC5B and tdTomato-CESA6, the GFP-SEC5B was transformed into plants expressing tdTomato-CESA6 in homozygous *prc1-1* background (Sampathkumar et al., 2013). The double-marked line of EXO70A1-GFP and tdTomato-CESA6 was generated by crossing. For plants co-expressing tdTomato-CESA6 and exocyst markers in the homozygous *xik-2* background, EXO70A1-GFP or GFP-SEC5B was crossed with tdTomato-CESA6 in homozygous *xik-2* and the F3 generation of *xik-2* homozygous plants expressing both markers were recovered. For plants co-expressing XIK-mCherry and YFP-CESA6, the XIK-mCherry construct was transformed into plants expressing YFP-CESA6 in homozygous *xik-2* and *prc1-1* mutant background.

Arabidopsis seeds were surface sterilized and stratified at 4°C for 3 d on half-strength Murashige and Skoog medium supplemented with 0.8% agar. For light growth, plants were grown under long-day lighting conditions (16 h light/8 h dark) at 21°C. For dark growth, plates were exposed to light for 4 h and then placed vertically and kept at 21°C in continuous darkness.

### Live-Cell Imaging

For most experiments, epidermal cells from the apical region of 3-d-old etiolated hypocotyls were imaged unless otherwise stated. Spinning-disk confocal microscopy (SDCM) was performed using a Yokogawa scanner unit (CSU-X1-A1; Hamamatsu Photonics) mounted on an Olympus IX-83 microscope, equipped with a 100X 1.45–numerical aperture (NA) UPlanSApo oil objective (Olympus) and an Andor iXon Ultra 897BV EMCCD camera (Andor Technology). YFP, GFP, and mCherry/tdTomato fluorescence were excited with 514-nm, 488-nm, and 561-nm laser lines and emission collected through 542/27-nm, 525/30-nm, and 607/36-nm filters, respectively. For measuring the abundance of plasma membrane (PM)-localized YFP-CESA6, time-lapse images were collected at the PM with a 2-s interval for 5 frames. For quantifying the abundance of cortical and subcortical CSC vesicles, z-series at 0.2 µm step sizes plus time-lapse with 1.6-s intervals for 10 frames were collected. For dual-wavelength imaging of GFP-SEC5B or EXO70A1-GFP with XIK-mCherry with SDCM, time-lapse images were collected at the PM focal plane with 1-s intervals for 2 min. For dual-wavelength imaging of GFP-SEC5B or EXO70A1-GFP with tdTomato-CESA6, time-lapse images were collected at the PM focal plane with 2-s intervals for 10 min. For dual-wavelength imaging of XIK-mCherry with YFP-CESA6, time-lapse images were collected at the PM focal plane with 3-s intervals for 10 min. For all dual-wavelength image acquisition, single-marked lines were tested initially to make sure there was no bleed through in each channel.

For imaging of single-marked lines of EXO70A1-GFP, SEC6-GFP and GFP-SEC5B, variable-angle epifluorescence microscopy (VAEM) was performed using a total internal reflection fluorescence (TIRF) illuminator on an IX-71 microscope (Olympus) equipped with a 150X 1.45–NA PlanApo TIRF objective (Olympus) and an EMCCD camera (ORCA-EM C9100-12; Hamamatsu Photonics). GFP fluorescence was excited with a 488-nm laser at 5% power and time-lapse images were collected at the PM focal plane with 0.5-s intervals for 1 min.

Fluorescence recovery after photobleaching (FRAP) experiments were performed as described previously (Zhang et al., 2019).

### Image Processing and Quantitative Analysis

Image processing and analysis were performed with Fiji Is Just ImageJ (Schindelin et al., 2012). The YFP-CESA6 related assays, including CSC density, delivery rate, cortical and subcortical CESA compartment density, and single CSC insertion assays, were performed as described previously (Zhang et al., 2019).

For exocyst subunit dynamics assay, the density of PM-localized exocyst foci was measured with the TrackMate plugin using the first frames of the time-lapse images. The Laplacian of Gaussians (LoG) detector was used and the estimated particle diameter was 4 pixels and the threshold set to 10. Stationary exocyst foci were tracked with TrackMate using the Simple LAP tracker with the maximal linking distance and gap-closing distance set to 1 pixel, which allowed the detection of foci that had no lateral motility. The frequency of stationary foci was calculated as the number of stationary foci divided by the measured area and elapsed time. For fluorescence intensity measurements of stationary foci at the PM, a fixed ROI of 3 × 3 pixels at the centroid of the foci was selected and analyzed over a time course. The first frame in which a new particle appears was set as 0 s. The fluorescence intensity of the ROI was measured from 6 frames (−3 s) prior to the first appearance of the particle to a few seconds after the foci fully disappeared. The lifetime of exocyst foci was measured by analysis of kymographs. Only a straight line in kymographs that could be tracked for at least 5 frames (>2 s) was measured.

For colocalization analysis of XIK-mCherry with GFP-SEC5B or EXO70A1-GFP, individual exocyst foci were tracked at the PM plane over a time course. Only stationary foci that could be tracked for at least 5 frames (> 4 s) were analyzed. For fluorescence intensity analysis, a fixed ROI of 3 × 3 pixels at the centroid of the foci was selected and the ROI was measured from 6 frames (−6 s) prior to the first appearance of the particle (0 s) to a few seconds after the foci fully disappeared. The same ROI in the corresponding XIK-mCherry channel was measured for fluorescence intensity at every time point. For normalization, the relative fluorescence intensity of GFP-SEC5B or EXO70A1-GFP was calculated as the fluorescence intensity of the ROI in each frame divided by the average fluorescence intensity of ROIs from the 6 frames prior to the first appearance of the particle, assuming that the fluorescence in those time points represents background signal. The relative fluorescence intensity of XIK-mCherry was calculated as the fluorescence intensity of the ROI in each frame divided by the average fluorescence intensity of ROIs from the last 6 frames in the time course when the foci were fully disappeared, assuming that the myosin signal in those time points represents random fluorescence.

The colocalization analysis of CESA6 with exocyst markers or myosin XIK was performed as previously described (Zhang et al., 2019). Individual CSC insertion events were tracked from erratic phase, through the pause phase, and into the beginning of the steady movement phase. Fluorescence intensities of ROIs in the corresponding exocyst or XIK channel were measured at each time point. The fluorescence intensities were normalized as described previously (Zhang et al., 2019).

### Yeast Two-Hybrid Assay

The cDNAs encoding *Arabidopsis* exocyst subunits were cloned into the prey vector pGADT7. The cDNA encoding the globular tail domain (GTD) of myosin XIK (amino acids 1055-1531) was amplified by PCR using primers GGAATTCCATATGATTTCGCCAACCAGCAGAACT and CCGGAATTCTTACGATGTACTGCCTTCTTTA and cloned into the bait vector pGBKT7. The bait and prey vectors were co-transformed into yeast AH109 strain and the positive clones were selected on SD medium lacking Trp, Leu and His and supplemented with 4 mM 3-amino-1,2,4-triazole. For negative controls, the XIK GTD bait vector was co-transformed with the empty pGADT7 vector and the empty bait vector was co-transformed with each prey vector containing the exocyst subunits.

### Protein Pull-Down Assay

The cDNA encoding the GTD (amino acids 1055-1531) of myosin XIK was amplified by PCR using primers TAGGATCCGGCGGTGGCGGTTCTATTTCGCCAACCAGCA and CGGAATTCGTTACGATGTACTGCCTTC and cloned into pGEX-4T-3 to generate an N-terminal GST fusion. The full-length cDNA for SEC5B was cloned into the pRSF-Duet vector to generate an N-terminal His6 fusion. Both constructs were transformed into *Escherichia coli* BL21 (DE3) and fusion protein expression induced with 0.5 mM isopropylthio-β-galactoside at 20°C for 3–6 h. The GST and GST-XIK fusion proteins were extracted with Pierce Immobilized Glutathione resin (Thermo Fisher Scientific) with the protein interaction buffer (20 mM Hepes-KOH pH7.2, 50 mM potassium acetate, 1 mM EDTA, 1 mM EGTA, 1 mM DTT, 5% glycerol, 0.5% Triton-100) as lysis and wash buffer. The His6-SEC5B fusion protein was extracted and purified with Ni-NTA His bind resin (Novagen). The freshly extracted resin-bound GST or GST-XIK proteins were mixed with purified His6-SEC5B protein in protein interaction buffer and incubated at 4°C for 2 h with shaking. The resin was washed 3 times with protein interaction buffer followed by boiling in SDS/PAGE sample buffer for 10 min, and then analyzed with SDS-PAGE.

### Cellulose Content Measurement

Five-d-old dark-grown hypocotyls were used for cellulose content assay. The alcohol-insoluble cell wall material was generated and hydrolyzed with acetic-nitric (AN) reagent or trifluoroacetic acid (TFA) based on the Updegraff method (1969) as described previously (Zhang et al., 2019). The insoluble fractions was then measured by a phenol-sulfuric colorimetric assay (Dubois et al., 1956) to determine the cellulose amount in the samples.

### Statistical Analysis

One-way ANOVA with Tukey’s post hoc tests were performed in SPSS (Version 25) to determine significance among different treatments. Two tailed Student’s t-tests were performed in Excel 15.32.

For statistical analysis by X-bar and S Control Chart, the upper control limit (UCL) was calculated in Excel 15.32 using the equation (Montgomery, 2009):

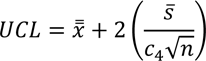

Any value higher than the UCL was considered as significantly different, with a P-value < 0.05.

### Accession Numbers

Sequence data from this article can be found in the Arabidopsis Genome Initiative under the following accession numbers: *Myosin XIK*, At5g20490; *Myosin XI1*, At1g17580; and *Myosin XI2*, At5g43900; *SEC3A*, At1g47550; *SEC5A*, At1g76850; *SEC5B,* At1g21170; *SEC6*, At1g71820; *SEC8*, At3g10380; *SEC10*, At5g12370; *SEC15A*, At3g56640; *SEC15B*, At4g02350; *EXO70A1*, At5g03540; *EXO84A*, At1g10385; *EXO84B*, At5g49830; *EXO84C*, At1g10180.

## Supplemental Data

The following materials are available in the online version of this article.

**Supplemental Figure 1.** Dynamic Behavior of EXO70A1-GFP Is Altered in *xik-1*.

**Supplemental Figure 2.** Dynamic Behavior of GFP-SEC5B Is Altered upon Inhibition of Myosin.

**Supplemental Figure 3.** Dynamic Behavior of Exocyst Subunits Is Altered upon Inhibition of Myosin and Actin.

**Supplemental Figure 4.** Myosin XIK Transiently Colocalizes with Stationary Foci of EXO70A1.

**Supplemental Figure 5.** EXO70A1 Transiently Colocalizes with CESA6 during the Vesicle Tethering Step of Secretion.

**Supplemental Movie 1.** A CSC Insertion Event at the PM.

**Supplemental Movie 2.** EXO70A1-GFP Distribution and Dynamic Behavior at the PM.

**Supplemental Movie 3.** Colocalization of GFP-SEC5B and XIK-mCherry Near the PM.

**Supplemental Movie 4.** Colocalization of GFP-SEC5B and XIK-mCherry during a Single SEC5B Arrival Event at the PM.

**Supplemental Movie 5.** Colocalization of GFP-SEC5B and tdTomato-CESA6 during a CSC Insertion Event at the PM.

## ACKNOWLEDGEMENTS

This work was supported by an award from the Office of Science at the US Department of Energy, Physical Biosciences Program, under contract number DE-FGO2-09ER15526 to C.J.S. We thank Nick Carpita and Anna Olek (Purdue) for assistance and access to equipment for cellulose determination. Cellulose analyses were supported by the Center for the Direct Catalytic Conversion of Biomass to Biofuels, an Energy Frontiers Research Center of the U.S. Department of Energy, Office of Science, Basic Energy Sciences (grant no. DE–SC0000997). We thank Valerian Dolja (Oregon State University) for providing the *myosin xi* triple knockout line and XIK-mCherry transgenic line, Ying Gu (Penn State University) for sharing the CESA6-YFP complementation line, David W. Ehrhardt (Carnegie Institution for Science) for the tdTomato-CESA6 line, and Viktor Žárský (Charles University) for the EXO70A1-GFP and SEC6-GFP lines. The authors are grateful to Hongbing Luo (Purdue) for excellent care and maintenance of plant materials.

## AUTHOR CONTRIBUTIONS

W.Z., L.H., C.Z., and C.J.S. designed the research. W.Z. and L.H. performed the experiments and data analysis. W.Z. and C.J.S. wrote the article.

